# O-GlcNAcylation and low glycolysis underpin Th2 polarization by dendritic cells

**DOI:** 10.64898/2026.02.03.703465

**Authors:** Leonard R. Pelgrom, Marjolein Quik, Jose J. Fernandez, Thiago A. Patente, Graham Heieis, Alexey A. Sergushichev, Jie Kang, Xiaobo Wang, Justine Dontaine, Marie-Sophie Fabre, Frank Otto, Alwin J. van der Ham, Lisa W. Bloemberg, Marion König, Beatrice M. F. Winkel, Rayman T.N. Tjokrodirijo, Arnoud H. de Ru, Rick Maizels, Meta Roestenberg, Tie Xia, Yan Shi, Olivier Lamiable, Luc Bertrand, Peter A. van Veelen, Maxim N. Artyomov, Cornelis H. Hokke, Bart Everts

## Abstract

Activation of dendritic cells (DCs) is dependent on rewiring of their cellular metabolism. However, the metabolic requirements for DCs to prime T helper 2 (Th2) responses are still poorly understood. Using unbiased transcriptomics and non-targeted metabolomics we find that helminth antigen-conditioned human DCs suppress glycolysis while increasing hexosamine biosynthesis to fuel protein O-GlcNAcylation. Functionally, glycolytic inhibition of DCs selectively enhanced, while blocking O-GlcNAcylation impaired, Th2-priming capacity. In helminth infection and allergic challenge, Th2 responses were also attenuated *in vivo* in mice with specific deletion of O-GlcNAc Transferase (OGT) in CD11c-expressing cells. Mechanistically, through proteomic analysis and functional validation, we identified O-GlcNAcylation as a critical negative regulator of immune synapse formation by controlling cytoskeletal organization via Fascin-1 and Zyxin, thereby dampening TCR signalling to promote Th2 polarization. Altogether we reveal a novel metabolic program in DCs that governs Th2 polarization, that could potentially be harnessed to treat type 2 mediated inflammatory diseases.

## Introduction

Type 2 immune responses are critical for host defense against parasitic helminths infection and for the regulation of tissue repair. Additionally, they can confer protection against metabolic and cardiovascular diseases, including type 2 diabetes. However, overzealous type 2 immunity can lead to allergy and support tumor growth^1^. The crucial involvement of type 2 immunity in disease signifies the importance of better understanding how they are initiated, to be able to devise novel approaches to target this response for clinical benefit.

The outcome of the immune responses depends centrally on the activation and polarization of CD4^+^ T cells. Dendritic cells (DCs) are the main professional antigen presenting cells required for T cell priming and differentiation towards different T helper (Th) subsets. There is a longstanding interest in delineating the cytokines and co-stimulatory molecules expressed by DCs that instruct CD4^+^ T cells to initiate a certain differentiation program^2^. These polarizing signals have been extensively characterized in the context of priming of Th1, Th17 and regulatory T (Treg) cell responses. Despite the necessity of DCs for initiating type 2 immunity, the exact signals that drive a CD4^+^ Th2 differentiation are far less well characterized. In part this stems from the fact that Th2-priming DCs do not appear to secrete polarizing cytokines analogous such as Interleukin (IL)-6, IL-12, IL-23, or Transforming Growth Factor (TGF)-β that contribute to Th1, Th17 or regulatory T cell (Treg) differentiation, respectively^3, 4^. Low expression of these polarizing cytokines, in particular IL-12 is considered a prerequisite for Th2 polarization by DCs, yet not sufficient. Over the years, ligands of the Notch receptor (Jagged 1 and 2) and the TNF receptor superfamily (OX40L) have been implicated in serving as possible membrane bound signals for promotion of Th2 polarization by DCs in certain contexts^5, 6, 7, 8^, although blockade of these pathways did not prevent Th2 priming *in vivo*^9, 10^. Additionally, it has been postulated that that reduced strength and duration of the cognate interaction between antigen-presenting DCs and CD4^+^ T cells favors differentiation towards the Th2 lineage^11^.

There is a growing appreciation that cellular metabolism is an additional aspect of DC function that can dictate their Th-priming bias^12, 13^. For instance, the production of the Th1-priming cytokine IL-12^14, 15, 16^ and Th17-priming IL-6 are dependent on glycolysis^17^. Similarly, tolerogenic DCs generated *in vitro* or isolated from tumors depend on fatty acid oxidation and mitochondrial respiration for the induction of Tregs^18, 19^,^20^. On the other hand, the metabolic properties of Th2-priming DCs and the metabolic programs that support this ability are still ill defined. House dust mite-stimulated murine DCs display an early induction of glycolysis *in vitro*^21^ and Irg1-driven itaconate production in DCs has been shown to suppress Th2-mediated airway inflammation, suggesting an important role for the canonical TCA cycle^22^. In addition, signaling through the metabolic sensor AMPK has been linked positively to Th2 polarization in the context of helminth infection^23^ while mTOR inhibition can favor human DCs for Th2 priming, together indicating Th2 priming by DC requires a metabolic shift towards catabolism^24^. Accordingly, evidence suggests that PPAR-g, a core regulator of fatty acid oxidation, is required for DC to drive Th2 responses^25^, however data is conflicting^26^, and whether PPARg acts through a metabolic mechanism in DCs remains to be shown. Altogether, a comprehensive understanding of how metabolism mechanistically governs Th2 polarization by DCs is currently still lacking.

Therefore, in the present study, we set out to characterize the metabolic properties of DC that allow them to prime Th2 cells. Using an *in vitro* model of Th2-priming by conditioning human-differentiated DCs with helminth antigens, we aimed to determine a metabolic signature of DCs with Th2-polarizing capacity through a combined experimental and computational pipeline of integrated metabolomics and transcriptomics datasets (CoMBI-T)^27^. We found that Th2-priming DCs can be distinguished from Th1- and Th17- priming DCs by their reduced glycolytic potential; a metabolic feature that in itself was sufficient to condition DCs for Th2 priming. Moreover, Th2-priming DCs displayed a distinct metabolic signature of UDP-GlcNAc synthesis through the hexosamine biosynthesis pathway, which forms the substrate for post-translational modification (O-GlcNAcylation) via the enzyme O-GlcNAc Transferase. Blocking protein O-GlcNAcylation, either pharmacologically or genetically, selectively reduced the Th2-priming capacity of these DCs *both in vitro* and *in vivo* in models of Type 2 immunity. Mechanistically, we find reduced glycolysis to limit IL-12 production and O-GlcNAcylation of Fascin-1 and Zyxin to suppress their cytoskeletal remodeling activity, resulting in reduced capacity of DCs to form strong immunological synapses with CD4^+^ T cells, thereby favoring Th2 polarization. Together, these findings define low glycolysis and protein O-GlcNAcylation as unique metabolic requirements for Th2 polarization by DCs and furthers our fundamental understanding of how type 2 immunity is initiated.

## Results

### Th2 priming human DCs are characterized by reduced glycolysis and enhanced amino-sugar metabolism

As a model to study the metabolic properties and requirements underlying Th2 polarization by DCs, we used the schistosome-derived glycoprotein omega-1 to condition human moDCs for Th2 priming (Th2-DCs). Omega-1 is a well-studied and potent inducer of Th2 responses both *in vitro* and *in vivo*^28, 29^. The transcriptome, generated by bulk RNA-sequencing, of these Th2-DCs was compared to those of unstimulated (immature DCs; iDCs), LPS-stimulated mature DCs (mDCs), LPS+Poly(I:C) stimulated Th1-priming (Th1-DCs) and zymosan-stimulated Th17-priming DCs (Th17-DCs; **Fig. 1a, b**). As previously reported^28^, Th2-DCs did not display a strong upregulation of costimulatory molecules, unlike Th1- and Th17-DCs (Extended Data **Fig. 1a**). Nonetheless, principle component analysis (PCA) of the transcriptome of Th2-DCs revealed it differed from both that of iDCs as well as of Th1-DCs and Th17-DCs (**Fig. 1c**; Extended Data **Fig. 1b**). Next, we generated transcriptional clusters with genes related to cellular metabolism probing for distinguishing signatures between the differently Th-polarizing DCs (**Fig. 1d**). Of the 13 generated clusters, the gene expression patterns of clusters 3 and 9 uniquely differentiated Th2-priming DCs from the other DCs; gene expression in cluster 3 was reduced and cluster 9 increased relative to the other DC populations. Pathway analysis was performed on metabolic genes in these clusters to elucidate which metabolic pathways were overrepresented. This revealed that in cluster 3 steroid biosynthesis had the highest rich factor, while glycolysis/gluconeogenesis was represented by the most genes (**Fig. 1e,f**). In cluster 9 lipid metabolism pathways were most significantly overrepresented although with a much lower rich factor than those represented in cluster 3 (Extended Data **Fig. 1c,d**).

**Fig. 1:**
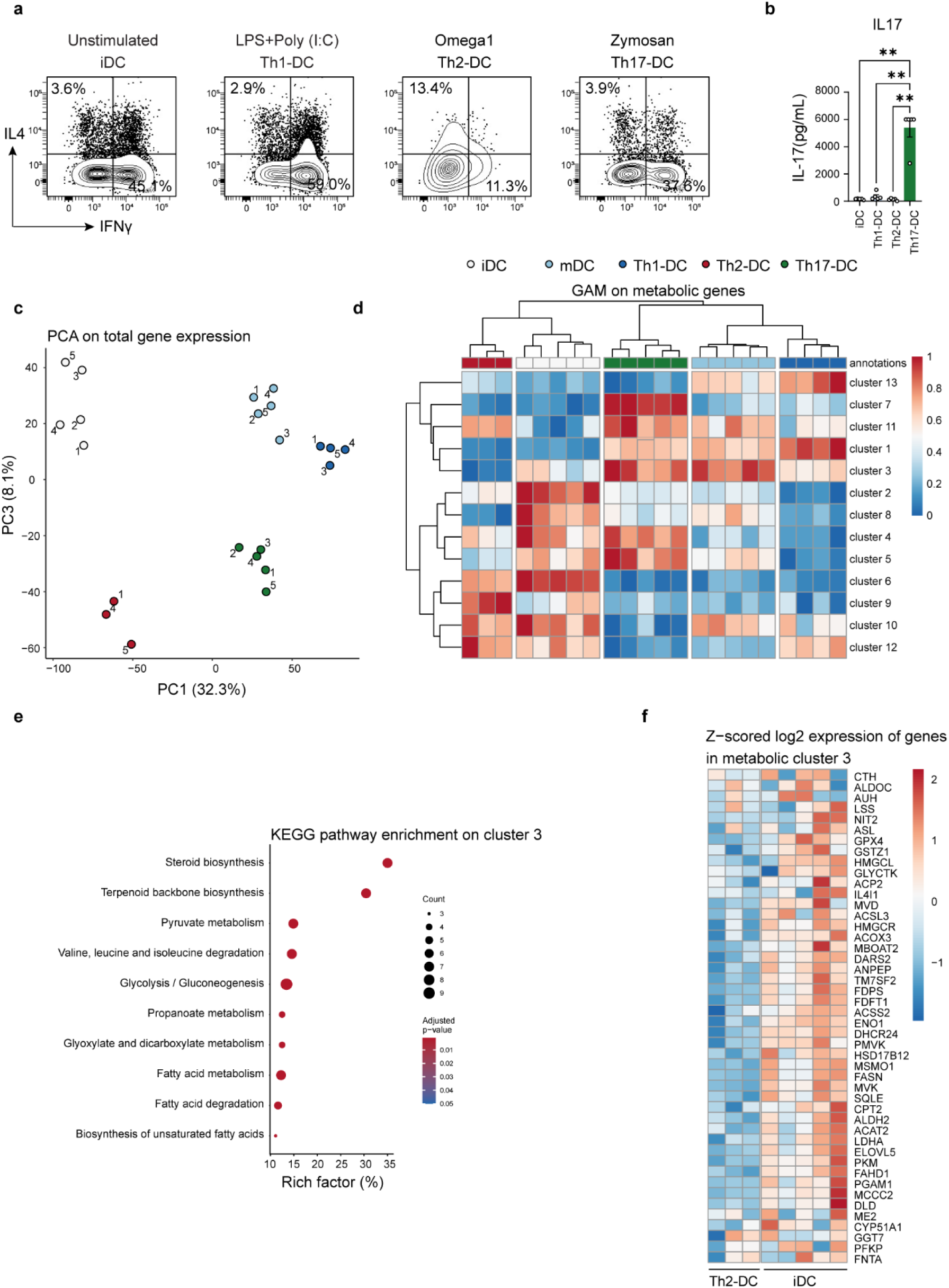
Metabolic characterization of Th2-priming DCs. Human moDCs were left untreated (iDC in white) or stimulated for 24h with either LPS+Poly (I:C) (Th1-DC in blue), zymosan (Th17-DC in green) or omega-1 (Th2-DC in red). **a**, Conditioned DCs were co-cultured with allogeneic naïve Th cells and 11 days later Th cell polarization was determined by flow cytometry. Representative flow plots of intracellular IFN-γ and IL-4 after a 6h stimulation with phorbol myristate acetate (PMA) and ionomycin. **b**, Conditioned-DCs were cocultured with memory Th cells and 5 days after DC-T cell coculture and IL-17 concentrations were determined by ELISA. **c,** Principal component analysis on total transcriptome of differently conditioned DCs. **d,** Metabolic genes were selected from the total transcriptome of differently conditioned DCs and grouped into 13 clusters optimized for maximal differentiation. **e**, KEGG pathways enrichment analysis of genes in cluster 3 and. **f**, Heatmap of genes within clusters 3. Genes and clusters with higher expression in a specific DC type compared to others are shown in red, while genes with lower expression are shown in blue. Data are based on 3 to 5 individual donors per condition. Statistical significance was determined by one-way ANOVA; \*\*\*\**p* < 0.0001.

To further define and compare functionally important nodes of metabolic rewiring in a non-targeted manner, we additionally performed unbiased metabolomics using flow injection analysis mass spectrometry (FIA-MS) in these cells^27^, yielding quantitative information on approximately 1200 MS features with 6000 putative metabolite annotations (Supplemental **Table 1**), and performed an integrated network analysis of these transcriptional and metabolomic data using GAM^30^. This integrated analysis corroborated our initial metabolic profiling based on transcriptomic data, as it identified suppressed glucose and fatty acid metabolism as main metabolic signatures of Th2-DCs in comparison to iDCs (**Fig. 2**). In addition, it revealed reduced TCA cycle activity. Interestingly, this analysis additionally uncovered a single significantly upregulated metabolic module linked to hexosamine biosynthesis, a pathway that fuels production of the amino sugar uridine diphosphate N-acetylglucosamine (UDP-GlcNAc), the substrate for a post-translational modification known as O-GlcNAcylation^31^. Together, this identifies a metabolic signature that is unique to Th2-priming human DCs, characterized by reduced glycolysis, fatty acid and TCA cycle metabolism, and increased UDP-GlcNAc metabolism.

**Fig. 2:**
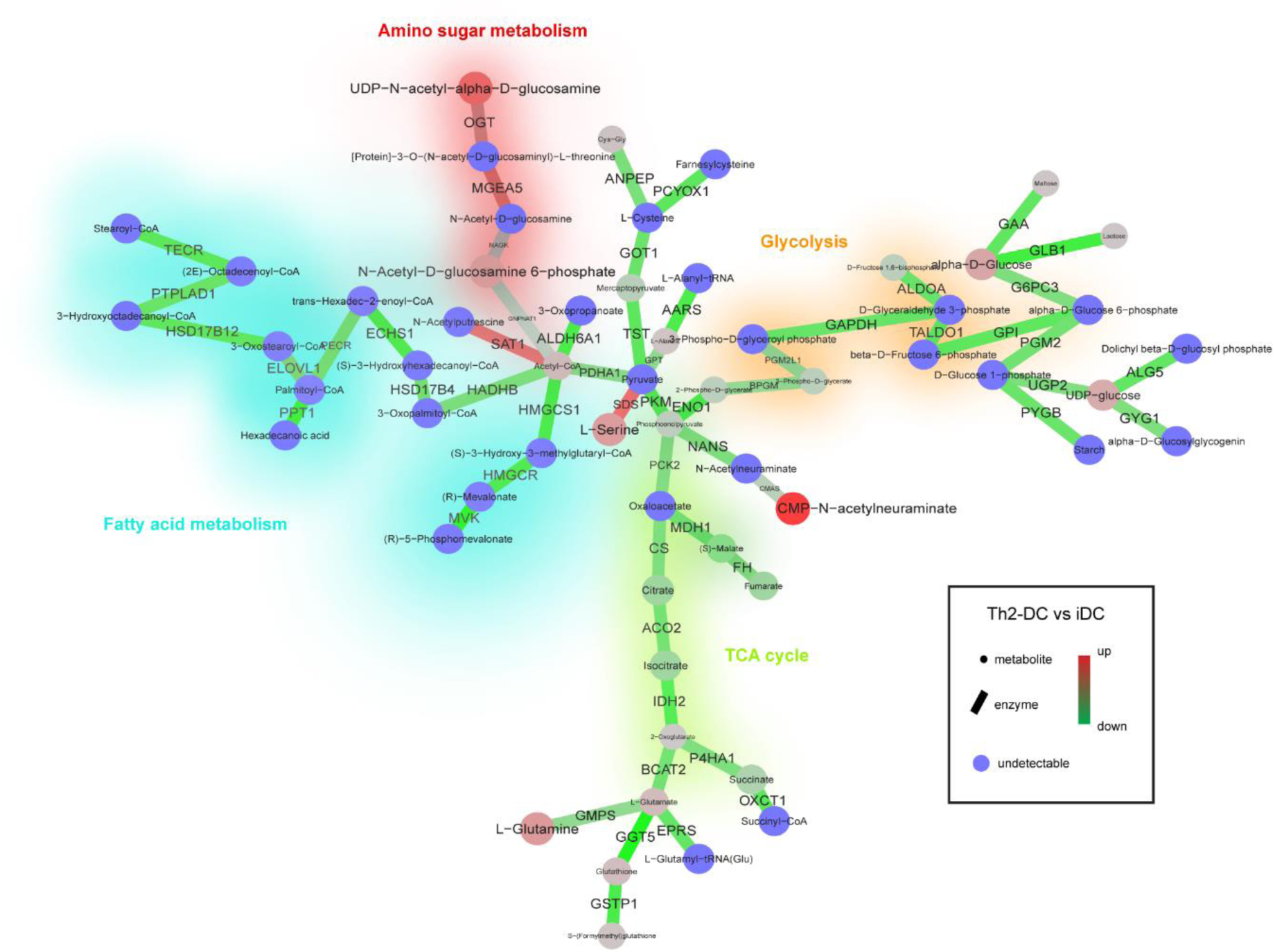
Metabolic network integration of Th2-priming DCs. Presentation of the most differentially regulated subnetworks within the global human metabolic network between Th2-DCs and iDCs that consists of enzymes and metabolites as determined through the CoMBI-T profiling pipeline based on integration of transcriptomic and metabolomic data determined by bulk RNA sequencing and flow injection analysis mass spectrometry (FIA-MS). Round nodes represent metabolites within the core regulatory network. Enzymes are represented by edges. Differential expression of enzymes or abundance of metabolites are indicated by red (higher in Th2-DCs) to green (higher in iDCs) color scale. Data are based on 3 to 5 individual donors per condition.

### Suppressed glycolysis underpins Th2-priming by DCs

To establish the biological relevance of these findings we first analyzed the glycolytic flux in differently conditioned DCs. Baseline rates of glycolysis were not significantly lower in Th2-DCs compared to iDCs as determined by extracellular flux (XF) analysis (**Fig. 3a**) and lactate release (**Fig. 3b**). Nonetheless, maximal glycolytic capacity (**Figure 3C**) and LPS-induced glycolytic reprogramming were impaired in Th2-DCs (Extended Data **Fig. 2a,b**). This metabolic phenotype was also observed when DCs were stimulated with Th2-priming house dust mite extract (Extended Data **Fig. 2a,b**). In contrast, Th1-DCs and Th17-DCs displayed strongly increased glycolytic rates and lactate release (**Fig. 3a,b**; Extended Data **Fig. 2c**). Th2-DCs did not compensate for a lower glycolytic flux by heightening the production of ATP through oxidative phosphorylation. In fact, consistent with the reduced expression of genes and metabolites related to the TCA cycle (**Fig. 2**), Th2-DCs displayed the lowest mitochondrial respiratory activity of all DCs, both in terms of baseline and maximal respiration as well as oxygen consumption linked to ATP synthesis (Extended Data **Fig. 2d,e**). This observation supports a low overall bioenergetic status of Th2-DCs.

**Fig. 3:**
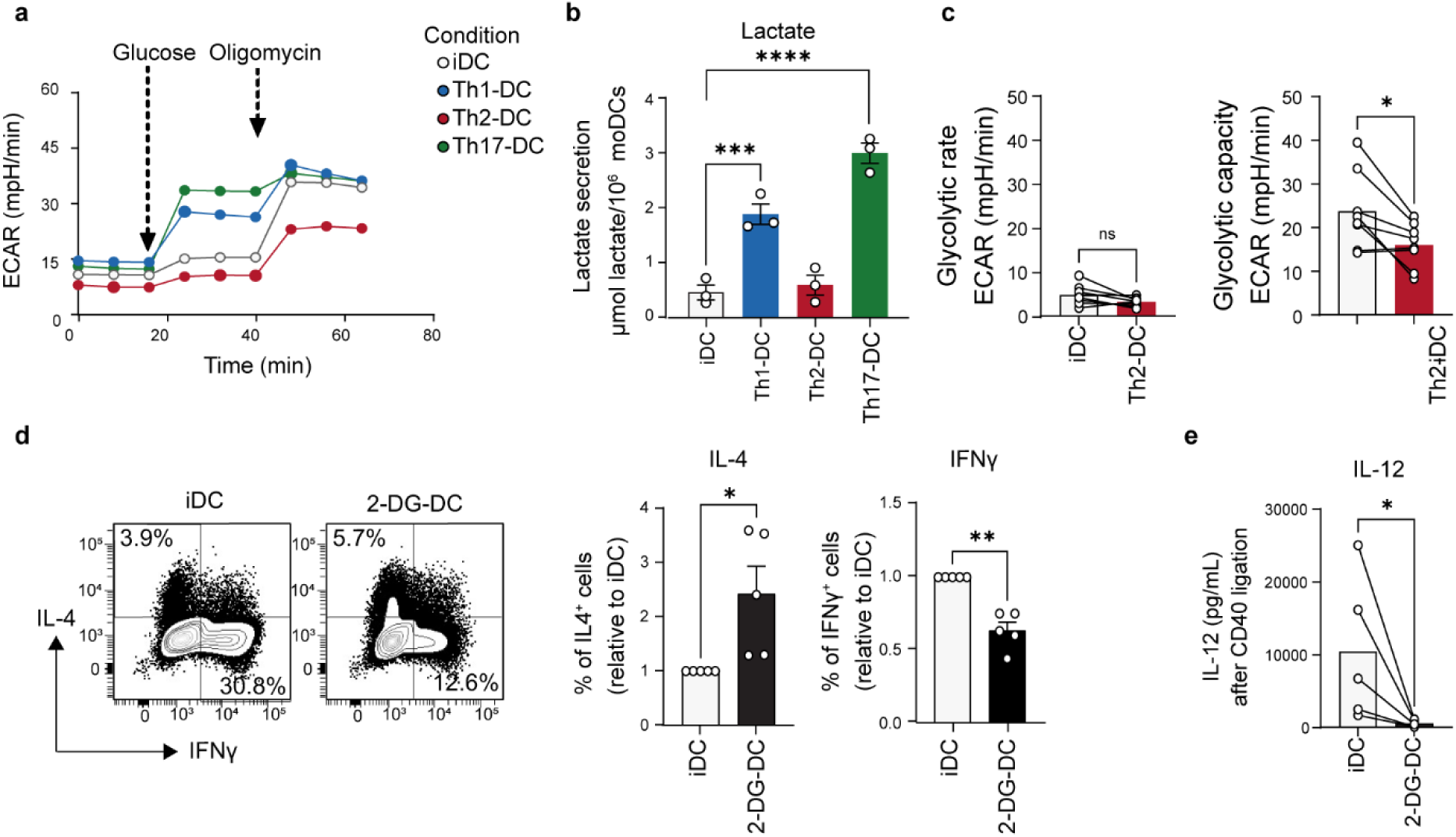
Suppressed glycolysis underpins Th2 priming by DCs. **a**, Glycolytic stress test in Seahorse XF analyzer which consists of sequential injections of glucose and oligomycin after which extracellular acidification rate (ECAR) is measured as a proxy for glycolysis-derived lactate. **b,** Supernatants from 24h conditioned human moDCs were collected for quantification of lactate release. **c**, Glycolytic rate is defined as the change in measurement before and after glucose injection. Glycolytic capacity is calculated as the difference between the baseline (pre-glucose injection) and the measurement following oligomycin injection. **d,** Human moDCs were left untreated or exposed for 24h to 2-deoxyglucose (2-DG-DC). Conditioned DCs were co-cultured with allogeneic naïve Th cells and 11 days later Th cell polarization was determined by flow cytometry. Representative flow plots of intracellular IFN-γ and IL-4 after a 6h stimulation with phorbol myristate acetate (PMA) and ionomycin. **e**, Conditioned DCs were co-cultured for 24h with a CD40L-expressing cell line after which supernatants were collected for determination of IL-12p70 secretion by ELISA. **a,** Data represent average of technical triplicates of one representative donor. **b-e**, Data points represent independent experiments of individual donors. Bars represent mean ± SEM; *p < 0.05, **p < 0.01, ***p < 0.001, ****p < 0.0001. Statistical significance was determined using one-way ANOVA with Dunnet post-hoc correction for multiple comparisons (**b**) and paired Student’s *t*-test (**c-e**).

The association between reduced glycolysis in Th2-DCs and their Th2-priming capacity led us to ask if lower glycolytic engagement is functionally linked to the Th2-priming capacity of these cells. Indeed, inhibiting glycolysis in iDC, using 2-deoxy-D-glucose (2-DG) promoted their ability to induce Th2 polarization (2-DG-DCs; **Fig. 3d)**, although not with the same potency as helminth antigens, confirming a role for metabolic rewiring in Th2 priming by DCs. A common characteristic and prerequisite of DCs for efficient Th2 priming is low IL-12 expression^28, 32^. Indeed, 2-DG-DCs secreted little IL-12 following CD40 ligation (**Fig. 3e**). Taken together, these data suggest that lower glycolytic activity in DCs supports their ability to differentiate Th2 cells by limiting IL-12 expression.

### O-GlcNAcylation is required for Th2 priming by DCs but not for Th1 or Th17 priming

The observation that glycolysis inhibition alone cannot fully recapitulate the full Th2-priming potential of helminth antigens, points to involvement of additional mechanisms. As our integrated omics analysis also identified hexosamine biosynthesis as a unique facet of Th2-DCs, we therefore aimed to better characterize and understand the role of O-GlcNAcylation. O-GlcNAcylation is regulated by two enzymes: O-GlcNAc transferase (OGT) and O-GlcNAcase (OGA; encoded by the *MGEA5* gene), that attach and remove O-GlcNAc moieties from proteins, respectively^31^ (**Fig. 4a**). Th2-DCs expressed higher mRNA levels of both *OGT* and *MGEA5* relative to iDCs (**Fig. 4b**; Extended Data **Fig. 3a**). Revisiting our metabolomics dataset, we additionally confirmed an increased intracellular concentration of the substrate UDP-GlcNAc in Th2-DCs relative to other Th-polarizing DCs, as well as its precursor N-acetyl-glucosamine-1/6-phosphate (**Fig. 4c,d).** However, expression of rate limiting enzymes in the hexosamine biosynthesis pathway, such as GFPT and GNA, was reduced in Th2-DCs relative to the other DCs (**Fig. 4b,e**). Instead, we observed increased mRNA expression of the enzyme NAGK, responsible for recycling of intracellular GlcNAc (**Fig. 4a,b,d**), pointing towards increased UDP-GlcNAc salvage rather than de novo synthesis in Th2-DCs. Consistent with this metabolic signature, we found increased levels of global O-GlcNAcylation as determined by flow cytometry (**Fig. 4f**). No major shifts in O-GlcNAcylation profile by immunoblot could be observed, indicating alterations in O-GlcNAcylation of a select number of proteins rather than global changes (Extended Data **Fig. 3b**). Taken together, these data show that in addition to suppressed glycolysis, Th2-priming DCs are metabolically wired to sequester UDP-GlcNAc, likely via the salvage pathway, for the maintenance of protein O-GlcNAcylation.

**Fig. 4:**
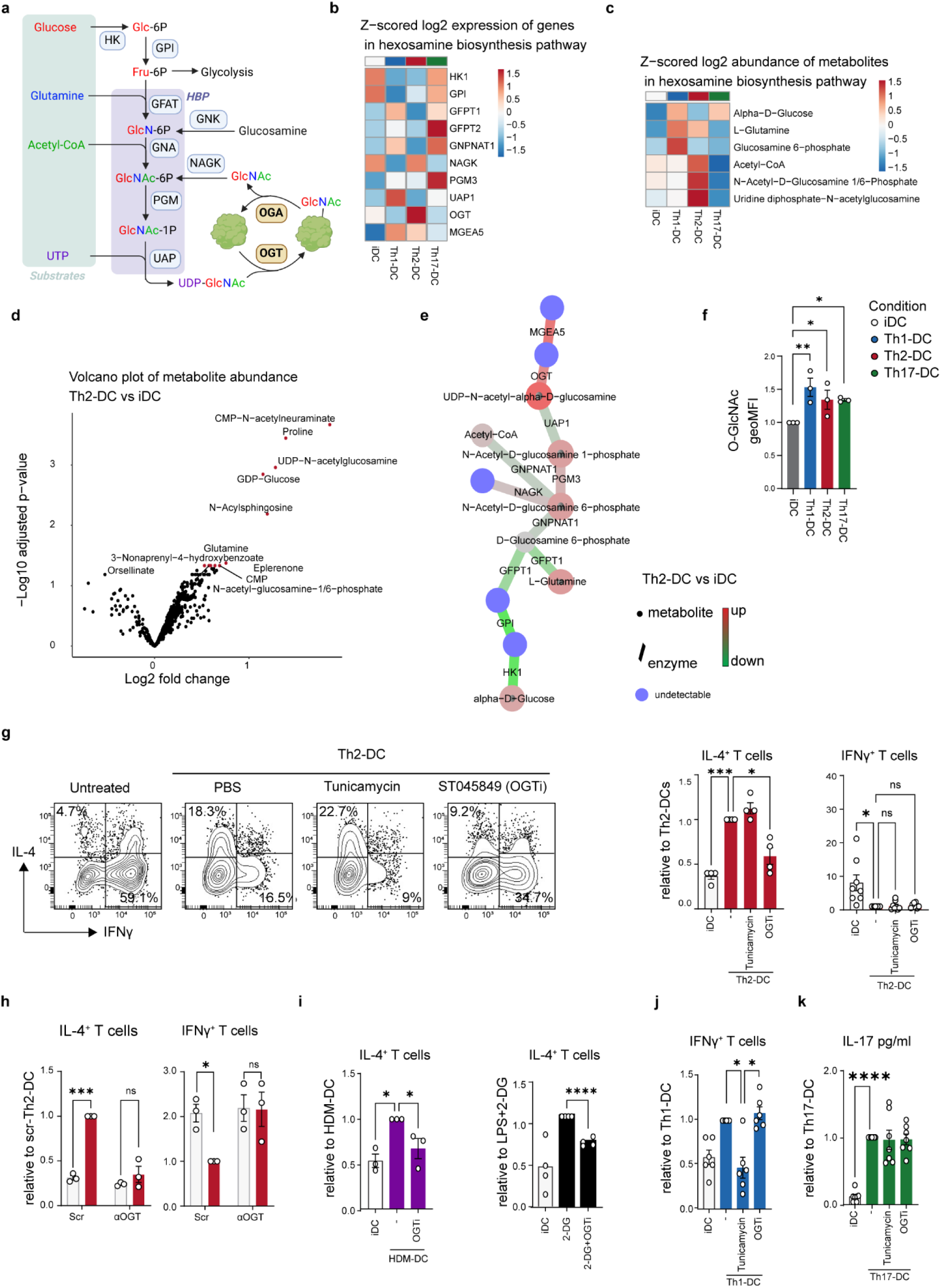
O-GlcNAcylation is required for Th2 priming by DCs. **a,** Schematic of the hexosamine biosynthetic pathway (HBP) and O-GlcNAcylation. **b,** Relative mRNA expression of 24h-conditioned human moDCs based on bulk RNA sequencing data (Figure 1). **c**, Relative intracellular metabolite abundance of 24h-conditioned human moDCs based on flow injection analysis mass spectrometry (FIA-MS; Figure 1). **d,** Volcano plot showing differences in intracelullar metabolite abundance between Th2-DCs and iDCs. Positive values corresponding to metabolites with increased abundance in Th2-DCs. Significantly different metabolites are highlighted in red. **e**, Integration of metabolomic and transcriptomic data sets using CoMBI-T profiling related to the HBP pathway (as described in Figure 2). Round nodes represent metabolites within the core regulatory network. Enzymes are represented by edges. Differential expression of enzymes or abundance of metabolites are indicated by red (higher in Th2-DCs) to green (higher in iDCs) color scale. **f,** Flow cytometry-based analysis of overall protein O-GlcNAcylation of conditioned human moDCs. **g, i-k,** Human moDCs were preincubated with inhibitors of O-GlcNAcylation (ST045849 – OGTi), or N-glycosylation (tunicamycin) for 30 minutes, before stimulation with **g**, omega-1, **i**, House dust mite (HDM) extract, 2-DG, **j**, LPS/Poly-IC or Zymosan, after which washed DCs were cocultured with (**g-j**) allogeneic naïve Th cells and 11 days later Th1/2 cell polarization was determined by intracellular cytokine staining or **k**, with memory Th cells and 5 days after DC-T cell coculture and IL-17 concentrations were determined by ELISA. (**G**) Representative flow plots are shown on the left. On the right % cytokine-producing T cells are shown relative to control conditioned-DCs. **h,** Differentiating moDCs were transduced with short interfering RNA against O-GlcNAc transferase (OGT) at day 4 and subsequently analyzed as in (**g**). Scrambled RNA was used as a control. Data points represent independent experiments of individual donors. Bars represent mean ± SEM; *p < 0.05, **p < 0.01, ***p < 0.001, ***p < 0.0001. Statistical significance was determined using one-way ANOVA with Dunnet post-hoc test (**f,g,i-k**) and Student’s *t*-test (**h**).

To test the functional importance of O-GlcNAcylation for Th2 priming by DCs *in vitro*, we inhibited OGT using the pharmacological inhibitor ST045849 (OGTi). This attenuated the capacity of Th2-DCs to prime IL-4-producing Th2 cells, without changes in IFN-γ or IL-17 production in these cultures (**Fig. 4g**; Extended Data **Fig. 3c**). Correspondingly, knockdown of OGT – using short interfering RNA – compromised the Th2-priming capacity of Th2-DCs as well as their ability to suppress Th1 responses (**Fig. 4h**). This dependency on O-GlcNAcylation was not restricted to helminth antigen-driven Th2 priming, as also HDM-conditioned and 2-DG-treated DCs were impaired in their Th2-priming ability when O-GlcNAcylation was inhibited (**Fig. 4i**). Importantly, OGT blockade in Th1- and Th17-DCs did not compromise their Th cell-polarizing ability (**Fig. 4j,k**). UDP-GlcNAc can also serve as a substrate for N- and O-linked glycosylation. However, the N-glycosylation inhibitor tunicamycin did not affect the Th2-priming capacity of Th2-DCs (**Fig. 4g**). Instead it specifically limited the ability of Th1-DCs to induce IFN-γ production by T cells (**Fig. 4j**). These effects were not secondary to altered survival of the DCs (Extended Data **Fig. 3d**). Taken together, these data show that the transfer of GlcNAc onto proteins by OGT is uniquely important for Th2 priming by DCs *in vitro*.

### O-GlcNAcylation is required for Th2 priming by DCs in the context of helminth infection and allergic asthma

To study the physiological relevance of these finding, we first assessed whether helminth antigens also altered O-GlcNAcylation in DCs *in vivo*. To this end, mice were challenged intradermally with fluorescently-labeled soluble egg antigens (SEA) from *S. mansoni*, of which omega-1 is a major constituent, after which Ag^+^ DCs were analyzed from the skin-draining lymph nodes (sdLN) for O-GlcNAcylation levels by flow cytometry (**Fig. 5a,b** and Extended Data **Fig. 4** for gating strategy). In line with our *in vitro* data, O-GlcNAcylation was elevated in Ag^+^ migratory KLF4-dependent CD11c^lo^CD11b^lo^ (DN)DCs, a skin DC subset that has been shown to be required for Th2 priming^33, 34^ (**Fig. 5c**). To further assess the *in vivo* role of DC O-GlcNAcylation in the T cell priming, *Ogt^flox/flox^* mice were crossed to *Itgax^cre^* mice to generate mice with a selective deficiency for OGT, and thereby O-GlcNAcylation, in CD11c^+^ cells. Splenic conventional (c)DCs from the conditional knockout mice (CD11c^Δ*Ogt*^) showed a near complete loss of OGT expression (Extended Data **Fig. 5a**), which resulted in a significant reduction in global protein O-GlcNAcylation in these cells (Extended Data **Fig. 5b**). KO efficiency and concomitant loss of O-GlcNAcylation was highest in male mice (Extended Data **Fig. 5a,c**), possibly due to the fact that the *Ogt* gene is located on the X-chromosome. Therefore, all further experiments were performed with male mice. Immunophenotyping of the spleen at steady-state revealed loss of O-GlcNAcylation had no effect on the frequency of major immune cell lineages or DC subsets (Extended Data **Fig. 5d,e**). Co-stimulatory and metabolic protein expression further remained comparable between splenic DCs from CD11c*^flox^* and CD11c^Δ*Ogt*^ mice (Extended Data **Fig. 5f**). However, CD11c^Δ*Ogt*^ mice harboured fewer cDCs within the lung and colon, but interestingly not the liver or small intestine (Extended Data **Fig. 5g**). CD11c^Δ*Ogt*^ mice were devoid of Langerhans cells in the sdLN, though migratory DNDCs were importantly present at a similar frequency to CD11c*^flox^* littermates (Extended Data **Fig. 5h**). Despite the loss of certain DC populations in some tissues, homeostatic T cell frequencies, activation status and cytokine producing potential were unaltered in CD11c^Δ*Ogt*^ compared to CD11c*^flox^* mice (Extended Data **Fig. 5i-k**). Therefore, in the absence of inflammation cDCs and T cells are largely unaffected by loss of O-GlcNAcylation. To further probe functional consequences of Ogt loss in DCs, we performed skin FITC painting to track DC antigen uptake, migration and maturation (**Fig. 5d**), however neither was compromised in CD11c^Δ*Ogt*^ mice (**Fig. 5e,f**).

**Fig. 5:**
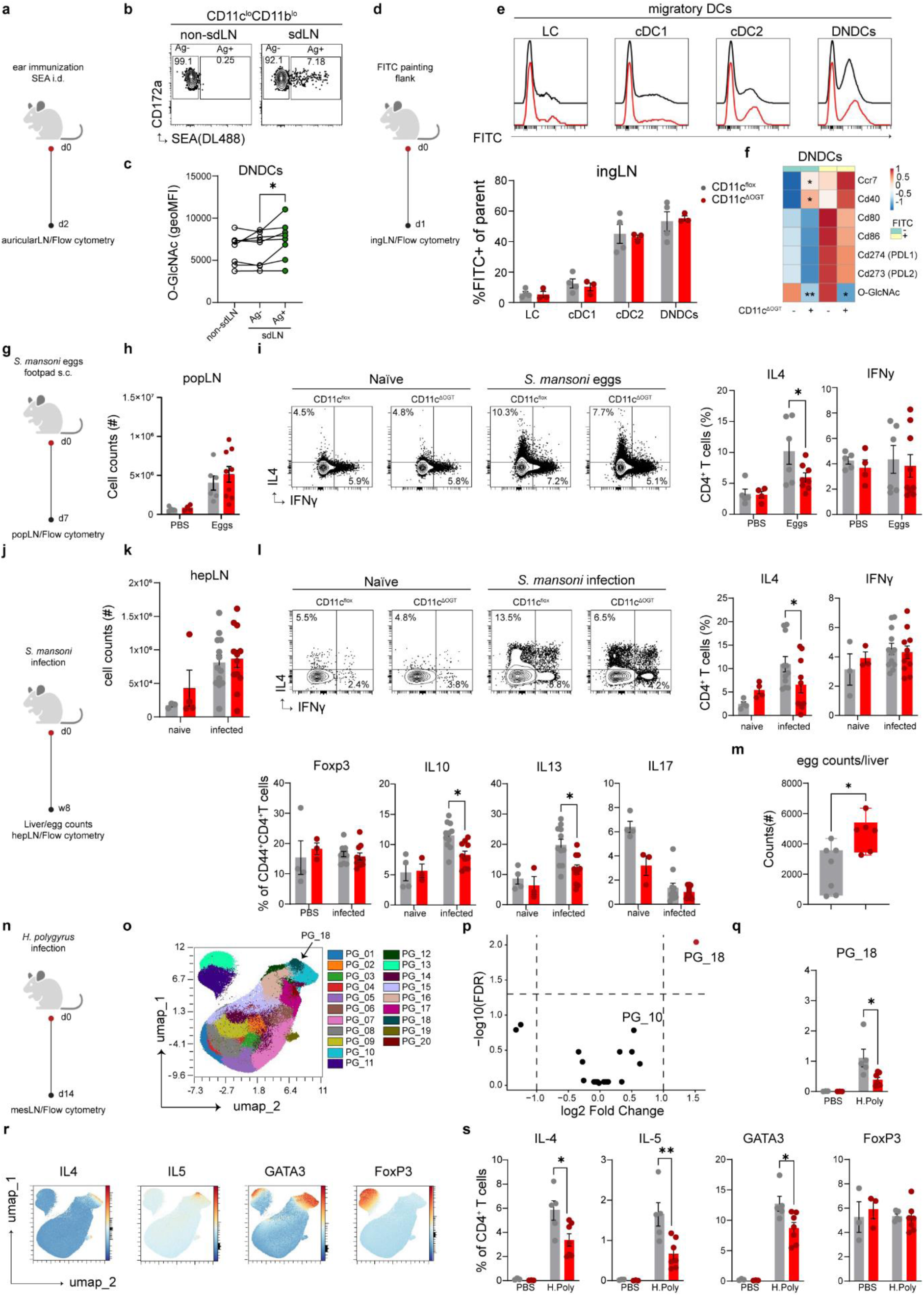
O-GlycNAcylation is required for Th2 priming by DCs *in vivo*. **a**, Mice were injected into ear pinnae with AF488-labelled SEA, after which 48h later protein **b,c**, O-GlcNAcylation levels were compared between in Ag negative and AF488-Ag^+^ CD11b^-^CD11c^-^CD24^-^double-negative DCs (CD11b^low^ = DNDCs) from draining LNs by flow cytometry. **d,** Migratory capacity of skin DCs was assessed by topical application of FITC paint on the flank, followed by analysis of FITC⁺ migratory DC subsets (migDCs) in the inguinal draining lymph node (LN) 1 day later. **e,** Flow cytometry analysis of FITC labelling in CD326^+^ Langerhans cells (LCs), XCR1^+^ cDC1, CD24^-^CD11b^+^ cDC2, and CD11b^-^CD11c^-^CD24^-^ double-negative DCs (DNDCs). **f,** Heatmap of log2 fold relative differences in expression of activation markers in splenic DCs from naïve CD11c^flox^ and CD11c^ΔOgt^ mice. Data are based on average of MFI of indicated markers from 3-4 mice per group. **g**, Mice were immunized in the footpad with *Schistosoma mansoni* eggs, and **h,** cell numbers and **i**, cytokine production by CD44⁺CD4⁺ T cells were analyzed in the popliteal LN 7 days post-immunization by ICS following PMA/Ionomycin restimulation. **j,** Mice were percutaneously infected with *S. mansoni* followed by the analysis of **k,** total cell numbers and **l,** cytokine production by CD44⁺CD4⁺ T cells in the hepatic LN by flow cytometry 8 weeks post infection. **m,** Quantification of liver egg burden 8 weeks after infection. **n,** Mice were orally infected with *H. polygyrus* L3 larvae, and mesenteric lymph nodes (mesLNs) were collected 14 days post-infection for flow cytometry analysis. **o,** PhenoGraph clustering overlaid on a UMAP representation generated from CD4⁺ T cell subsets. **p**, EdgeR analysis showing differential PhenoGraph cluster abundance between infected mice from CD11c^Δ*Ogt*^ and CD11c^flox^ littermates, of which **q**, frequencies of significantly reduced cluster PG_18 (highlighted by the arrow in (**o**)) in CD11c^Δ*Ogt*^ mice is displayed. **r**, Continuous color scales for IL-4, IL-5, GATA3, and Foxp3 expression across the PhenoGraph-clustered UMAP. **s**, The percentage of CD4⁺ T cells expressing IL-4, IL-5, GATA3, and Foxp3. Each symbol represents an individual mouse from (**c,e,f,m**) 1 of 2 independent experiments, (**h,i,k,l**) 2 independent experiments combined or (**o-s**) one experiment. Data are shown as mean ± SEM. Statistical significance was determined using Student’s t-test or two-way ANOVA with Tukey post-hoc test; \**p* < 0.05, \*\**p* < 0.01.

Having shown that DCs are largely unaffected by the loss of OGT in steady-state, we tested our hypothesis that O-GlcNAcylation in DCs is a required for *in vivo* Th2 priming. Mice were immunized in the footpad with eggs derived from the parasitic worm *Schistosoma mansoni,* which elicit a strong local type 2 immune response (**Fig. 5g**). 7 days after immunization, CD11c^Δ*Ogt*^ mice displayed similar increase in cellularity, but impaired Th2 induction in the draining popliteal LN, without a shift towards a Th1 response (**Fig. 5h,i**). To extend these findings to a natural infection, mice were infected with *S. mansoni*. At 8 weeks post-infection, corresponding with the peak of the Th2 response in the liver due to sexual maturation and egg production, we assessed the CD4^+^ T cell response from the liver-draining hepatic LN. Overall cell counts were again comparable between genotypes, and no differences were observed in IFN-γ-producing Th1 or Foxp3^+^ Tregs (**Fig. 5j,l**). Similar to egg-injection, natural infection induced a weaker Th2 response in CD11c^Δ*Ogt*^ animals compared to CD11c*^flox^*. In agreement with reduced Th2-driven immunity, parasite fecundity was greater in mice when DCs lacked OGT, as seen from increased egg counts in the liver (**Fig. 5m**). We further substantiated these findings in an alternative model of intestinal helminth infection using *Heligmosomoides polygryus* (**Fig. 5n-s**), as well as in house dust mite induced allergic inflammation (Extended Data **Fig. 6a-d**). In both cases, CD11c^Δ*Ogt*^ mice showed a similar selective impairment in their ability to prime a type 2 immune response compared to CD11c*^flox^* mice. Importantly, when CD11c^Δ*Ogt*^ mice were immunized with the TLR9 ligand CpG-B in combination with a synthetic 43-mer-long peptide from the human papillomavirus (HPV) E7 protein that contains the immunodominant and H-2Db-restricted epitope RAHYNIVTF (E7-SLP), a combination that predominantly activates cDC1s and requires cross-presentation for priming of CD8^+^ T cells^35^, no defect in the CD8^+^ T cell response could be observed (Extended Data **Fig. 6e-h**). Together this establishes that O-GlcNAcylation in DCs contribute to efficient CD4^+^ T cell priming, and is specifically required for Th2 responses *in vivo*.

### O-GlcNAcylation of Zyxin and Fascin-1 in DCs dampens strength of immunological synapse formation to favor Th2 polarization

Our data establish a clear role for O-GlcNAcylation in DC to efficiently prime Th2 cells both *in vitro* and *in vivo*. Thus we returned to our *in vitro* system to further interrogate the molecular mechanisms through which O-GlcNAcylation controls the Th2-priming capacity of DCs. Silencing of OGT had no effect on IL-12 production, contrasting glycolytic inhibition, suggesting an IL-12 independent mechanism (Extended Data **Fig. 7a**). O-GlcNAcylation, similar to phosphorylation, alters the functional properties proteins by covalently linking to serine-threonine residues. However, unlike kinases, OGT is highly promiscuous and modifies thousands of proteins^36^. We therefore compared the O-GlcNAcylome of Th2-DCs and unstimulated DCs to identify differentially O-GlcNAcylated proteins, and thereby provide leads for molecular targets through which O-GlcNAcylation supports Th2 priming. To this end, we applied a proteomics approach, based on click chemistry-dependent tagging and subsequent immunoprecipitation of O-GlcNAcylated proteins, followed by LC-MS/MS analysis for protein identification and quantification^37^ (Extended Data **Fig. 7b**).

Approximately 700 O-GlcNAcylated proteins were identified (Suppplemental **Table 2**). Of these, 30 were differentially O-GlcNAcylated, with 7 showing significantly reduced and 23 showing increased O-GlcNAcylation in Th2-DCs versus unstimulated DCs (**Fig. 6a**). Many of these 30 hits were ribosomal proteins that are well known targets of O-GlcNAcylation^38^. After excluding ribosomal hits, we notably identified Zyxin and Fascin-1, both actin remodeling proteins, that were increased in O-GlcNAcylation in Th2-DCs. (**Fig. 6b,c**). While O-GlcNAcylation of Zyxin has been identified and impairs its cytoskeletal remodeling activity^39^, O-GlcNAcylation of Fascin-1 has not been reported. Fascin-1 in DCs has been shown to mediate actin bundling required for immunological synapse formation with T cells^40^. Taken together, these observations led us to postulate a role for O-GlcNAcylation in controlling the strength of TCR signals provided to cognate CD4^+^ T cells, in order to instruct Th2 differentiation^11^.We hypothesized that increased O-GlcNAcylation of Zyxin and Fascin-1 would impair their cytoskeletal remodeling properties, thereby rendering DCs less capable of forming strong synapses with T cells allowing for Th2 differentiation.

**Fig. 6:**
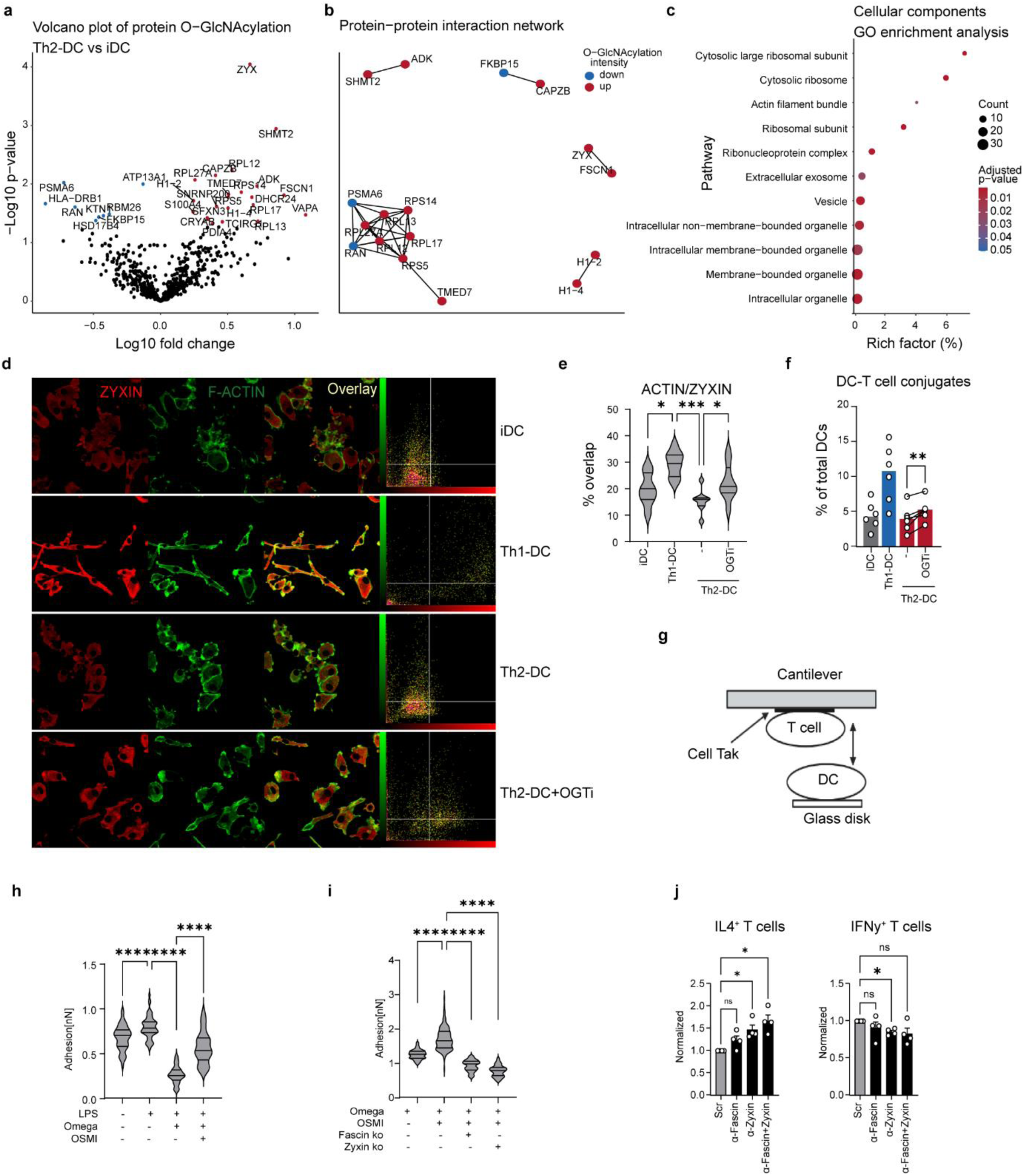
O-GlcNAcylation of Zyxin and Fascin-1 promotes Th2 priming by DCs by limiting immunological synapse strength. (**A**) MS/MS based proteomic analysis of O-GlcNAcylated proteins in differently conditioned human moDCs. Volcano plot showing differential protein O-GlcNAcylation in Th2-DCs compared with iDCs. (**B**) Protein–protein interaction network of proteins exhibiting increased or decreased O-GlcNAcylation in Th2-DCs. (**C**) Gene Ontology (GO) enrichment analysis of cellular component terms associated with differentially O-GlcNAcylated proteins in Th2-DCs. (**D**) Confocal microscopy of F-Actin (green), Zyxin (red), and merged images of moDCs left untreated or stimulated for 24 h with LPS, omega1, or omega1 preincubated with the O-GlcNAcylation inhibitor ST045849 (OGTi) for 30 minutes. (**E**) quantification of the percentage of overlap between F-Actin and ZYXIN in individual cells. (**F**) Conditioned human moDCs were co-cultured with allogenic naïve Th cells for 4h. DC-T cell conjugates were identified as CD1a^+^CD4^+^ dual-positive events and quantified as the percentage of total DCs. (**G**) A schematic diagram for Atomic Force Microscopy (AFM)-based single-cell force spectroscopy (SCFS) assay setup. (**H**) DC2.4 cells were stimulated with LPS or omega1 with or without pre-incubated with OGTi prior to stimulation. Adhesion was quantified as in (G). (**I**) WT, Fascin ko or Zxyin ko DC2.4 cells were stimulated with LPS or omega1 with or without pre-incubated with OGTi prior to stimulation. Adhesion assay following knockout as in (G) (**J**) siRNA mediated knockdown of Fascin-1 or Zyxin was performed on day 4 of moDCs differentiation; scrambled siRNA served as control. On day 6, DCs were co-cultured with naïve allogeneic T cells for 11 days. Intracellular IFN-γ and IL-4 production was assessed by flow cytometry after 6h of PMA/ionomycin stimulation. Percentages of Th cells expressing IFN-γ or IL-4 are shown relative to untreated scrambled control. (**A-C**) Analysis based on data from 4 donors. One representative of (**D**) 4 or (**H,I**) 2 independent experiments is shown. (**F,J**) Datapoints represent individual donors from independent experiments. Data are shown as mean ± SEM. Statistical significance was determined by (E) paired Student’s t-test or (G-I) using two-way ANOVA with Tukey post-hoc test; \**p* < 0.05, \*\**p* < 0.01, \*\*\**p* < 0.001, \*\*\*\**p* < 0.0001.

To test our hypothesis we first assessed Zyxin dynamics within DCs in response to omega-1 or LPS by confocal microscopy. Indeed, colocalization of Zyxin with the Actin cytoskeleton was lower in omega-1- compared to LPS-treated DCs and this was increased in the presence of OGT inhibitor OSMI (**Fig. 6d,e**). Further support for this model came from the observation that, in human DC-CD4^+^ T cell co-cultures, the frequency of DC-T cell doublets was low in Th2-priming cultures, as detected by flow cytometry. Furthermore, and OGT inhibition increased the doublet frequency (**Fig. 6f**). To directly measure the adhesion force between individual pairs of DCs and T cells we applied Atomic Force Microscopy (AFM)-based single-cell force spectroscopy (SCFS) using murine OVA-pulsed DCs and antigen specific OTII CD4^+^ T cells^41^ (**Fig. 6g**). In line with the human DC data, we found omega-1 to reduce adhesion strength between DCs and T cells and to antagonize LPS-induced interaction strength in an Ogt-dependent manner (**Fig. 6h**; Extended Data **Fig. 7c**). To directly link this to Zyxin and Fascin-1, CRISPR-Cas9 was applied to create Fascin-1- and Zyxin-deficient DCs (Extended Data **Fig. 7d**). Omega-1-treated DCs, deficient for either Fascin-1 or Zyxin, failed to increase adhesion force following Ogt inhibition, connecting O-GlcNAcylation of Fascin-1 and Zyxin to synapse strength (**Fig. 6i**). Finally, we tested whether silencing of *ZYX* and *FASCN1* could phenocopy omega-1-induced Th2 polarization by human moDCs. Silencing *ZYX* and *FASCN1* in unstimulated DCs was sufficient to render these cells capable of priming a Th2 response (**Figure 6J; S7E**). Together, these findings provide evidence for a key role for O-GlcNAcylation of Zyxin and Fascin-1 in underpinning Th2 priming by DCs by reducing DC-CD4^+^ T cell interaction strength.

### cDC2s from helminth infection individuals are characterized by increased protein O-GlcNAcylation

Finally, to study the human *in vivo* relevance of our findings, we examined whether protein O-GlcNAcylation was affected in peripheral blood DCs of participants taking part in a controlled human helminth infection (CHI) study using *Necator americanus^42^* (**Fig. 7a**). Eight weeks post infection, around the peak of the type 2 immune response as determined by eosinophilia (**Fig. 7b**), protein O-GlcNAcylation was detected by flow cytometry. O-GlcNAcylation was increased in type 2 conventional DCs (cDC2s) - the subset that is classically associated with the priming of Th cell responses^43^ - but not plasmacytoid DCs or cDC1s (**Figure 7c**). This increase shows that enhanced protein O-GlcNAcylation is also a characteristic of DCs during an active type 2 immune response in humans.

**Fig. 7:**
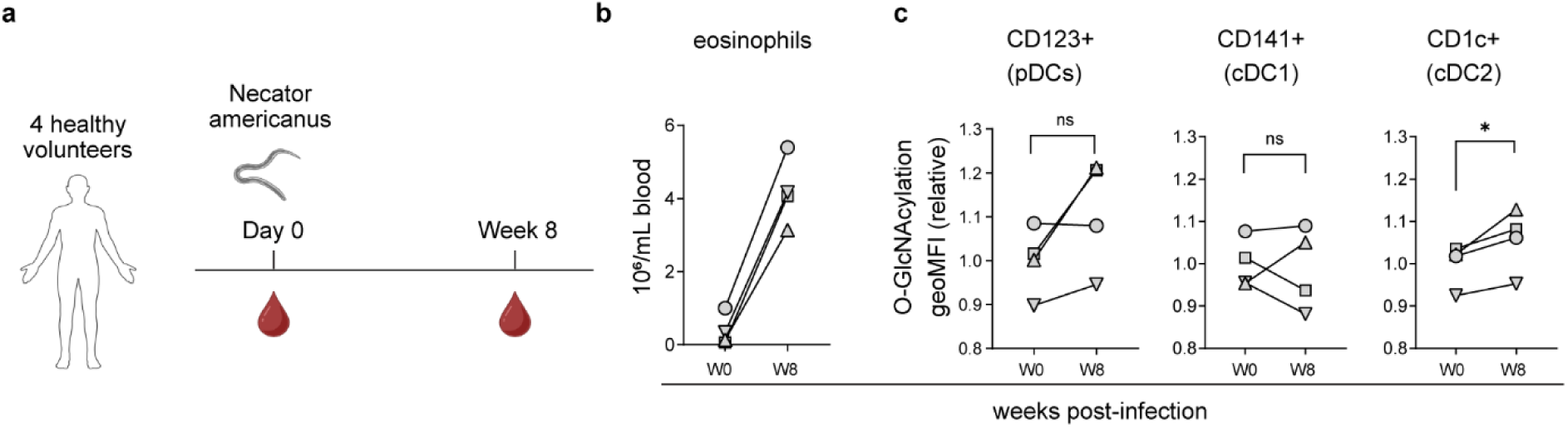
cDC2s from helminth infection individuals are characterized by increased O-GlcNAcylation. (**A**) 4 healthy volunteers were infected with the hookworm *Necator americanus* for 8 weeks in a controlled human infection (CHI) study. (**B**) Induction of type 2 immune responses was determined by quantification of eosinophilia in blood using flow cytometry (**C**) Global protein O-GlcNAcylation in peripheral blood CD123^+^ plasmacytoid DCs (pDCs), CD141^+^ type 1 conventional DCs (cDC1s) and CD1c^+^ (cDC2s) was assessed using flow cytometry. Data is shown as mean; each symbol represent a single volunteer; *p < 0.05. Statistical significance was determined by paired student’s T test.

## Discussion

Given the paucity in our understanding of the role of cellular metabolism in Th2 polarization by DCs, we aimed in the present study to identify metabolic properties that are unique to DCs that induce Th2 polarization and to determine the functional importance of those metabolic programs in DCs in priming this response. We find that Th2-priming DCs have a unique metabolic signature characterized by low glycolytic activity, a characteristic that we find to support the Th2-polarizing ability of DCs. Moreover, we found Th2-priming DCs to display a distinct metabolic signature of UDP-GlcNAc accumulation and enhanced O-GlcNAcylation. Blocking protein O-GlcNAcylation, either pharmacologically or genetically, selectively reduced the Th2-priming capacity of these DCs both *in vitro* and *in vivo* in models of type 2 immunity. Mechanistically, we provide evidence for reduced glycolysis to limit IL-12 production and for O-GlcNAcylation of Fascin-1 and Zyxin to suppress their cytoskeletal remodeling activity, resulting in reduced capacity of DCs to form strong immunological synapses with CD4^+^ T cells, which together favor Th2 polarization.

As opposed to DCs that prime Th1 or Th17 responses, Th2-priming DCs are commonly characterized by a muted activation phenotype and low expression of pro-inflammatory cytokines. Particularly, low IL-12 expression and a low antigen presentation have been linked to their ability to prime Th2 responses^44^. Induction of aerobic glycolysis has been shown to be associated with and required for acquisition immunogenic DC activation and production of Th1-polarizing signals such as IL-12^15, 28, 45^. Mechanistically, this has been linked to glycolysis-driven fatty acid synthesis that allows for ER/Golgi expansion in DCs after TLR ligation^14^, which is thought necessary for optimal expression of costimulatory molecules and T cell-polarizing cytokines. Therefore, the impaired glycolytic potential of Th2-DCs we observed in our work may ‘lock’ them into a muted activation phenotype as a prerequisite for efficient Th2 priming. Indeed, inhibition of glycolysis prevented IL-12 secretion by DCs following CD40 ligation and was sufficient to enhance their Th2 priming capacity. Although we provide evidence that both helminth antigens and allergens induce a glycolytically low DC phenotype, recent work identified an increased glycolytic signature in Mgl2^+^ cDC2s exposed to spores from *Aspergillus fumigatus*, that promote fungal allergic responses in the lung ^46^. Allergic inflammation induced by *A. fumigatus* also has a strong type 17 immune response component, which is known to be supported by CLR-Syk signaling-driven glycolysis in DCs^21, 47^. To what extent this glycolytic shift specifically underpins Th2-associated allergic inflammation in this model was not addressed.

We additionally identified elevated levels of HBP metabolites and O-GlcNAcylation as a defining metabolic signature of Th2-priming DCs. We specifically found increased expression of HBP enzymes and abundance of metabolites linked to GlcNAc recycling and O-GlcNAcylation, while those involved in *de novo* UDP-GlcNAc synthesis were not altered or reduced. The reduction in glycolysis observed in Th2-DCs is consistent with the preference for GlcNAc recycling, as glucose is a key substrate for de novo UDP-GlcNAc synthesis. In T cells, liver, neuronal and lung cancer cell lines, this inverse relation has also been reported, in which increases in OGT expression and protein O-GlcNAcylation are observed after glucose deprivation or glycolysis inhibition^48, 49, 50, 51^. It is therefore conceivable that in Th2-DCs, active suppression of glycolysis may be contributing to Th2 priming by DCs, not only as a prerequisite for Th2 induction by limiting their IL-12 expression and restricting overall maturation, but also by increasing UDP-GlcNAc utilization and O-GlcNAcylation.

The investigation of how O-GlcNAcylation effects the function of immune cells is still in its infancy, but is gaining traction^52^. Thus far, a single study has probed the role of O-GlcNAcylation in DC biology, revealing inhibition of O-GlcNAcylation affects human moDC differentiation from monocytes^53^. Yet, functional consequences or underlying molecular mechanisms were not assessed in this study. We here provide evidence that O-GlcNAcylation in differentiated DCs is required for Th2 priming by limiting strength of immunological synapse formation with T cells by targeting cytoskeletal remodeling proteins Zyxin and Fascin-1. Zyxin interacts with α-actinin and vasodilator-stimulated phosphoprotein to regulate actin dynamics and actin-membrane connections. Fascin-1 cross-links actin filaments (F-actin) into bundles that support tubular membrane protrusions including filopodia and stereocilia. In contrast to Zyxin, the role of Fascin-1 has been studied in DC biology before^54^. Fascin-1 plays a critical role in enhancing DC motility through actin cytoskeleton reorganization^55^. In addition, reduced Fascin-1 expression hampers cytoskeleton rearrangement in DCs/Langerhans cells, resulting in decreased T-cell activation^56, 57^, as a consequence of disrupted actin dynamics that control contact duration and priming efficiency at the immunological synapse with T cells^58^. Interestingly, O-GlcNAcylation of Zyxin was recently reported to promote nuclear translocation, thereby relieving it from its cytoskeletal remodeling properties^39^. On the other hand, although Fascin-1 harbors multiple putative O-GlcNAcylation sites (from online database O-GlcNAcPRED-DL), O-GlcNAcylation of Fascin-1 has not been documented, nor how its function is affected by it. Here we report that both Zyxin and Fascin-1 are O-GlcNAcylated in DCs upon helminth antigen stimulation. We accordingly observed that DC-T cell interaction strength is reduced upon helminth antigen exposure. Interaction strength was restored by inhibiting O-GlcNAcylation, but notably restoration required the expression of Zyxin and Fascin-1. Altogether our observations, paired with those previously reported, support a model in which enhanced O-GlcNAcylation specifically on Zyxin or Fascin-1 lowers the ability of DCs to form stable immunological synapses with T cells to favor Th2 differentiation.

Our findings align with several other studies in which lowering antigen dose, targeted mutations reducing TCR signal transduction and restricted T cell repertoires have been linked to driving Th2 differentiation as a consequence of reduced TCR signaling through the early promotion of GATA3 expression^11, 44, 59^. Conceptually, it seems unlikely that peptides from all allergens would drive low-strength, low-affinity interactions between DCs and CD4^+^ T cells, given their great diversity and often high availability. Our work suggests that this is indeed not an intrinsic property of the allergen but rather controlled by the antigen-presenting DCs that are actively programmed to limit the strength of interaction with CD4^+^ T cells independent from antigen dose or peptide affinity. There has been a longstanding quest for identifying immunological Th2 polarizing signals that can uniformly distinguish Th2-priming DCs from other DCs. Lack of identification of such signals, originally led to a ‘default concept’^60, 61^, which postulates that the absence of other Th-polarizing signals leads - by default - to a Th2 outcome^3^. Our work clearly argues against this passive ‘default concept’ for Th2 priming as it provides further support for immunological synapse strength as an active immunological signal governing Th2 polarization. Importantly, we have delineated a molecular mechanism through which the immunological synapse strength is controlled in DCs.

Our *in vivo* studies suggest that O-GlcNAcylation does not appear to play a major role in DC homeostasis in most analyzed tissues, or migration following inflammatory challenge. Correspondingly, at steady state, splenic T cell populations were not affected by loss of OGT in DCs, pointing towards no gross homeostatic functional consequences from the impairment of O-GlcNAcylation in DC populations. In contrast, intrinsic deletion of OGT in T cells blocks their activation, expansion and survival^62^. Similarly, in parallel work we observe disruption of O-GlcNAcylation in macrophages enforces a pro-inflammatory phenotype associated with widespread loss of tissue resident populations, and an associated systemic increase in homeostatic T cell activation^63^. Possibly, the known importance for O-GlcNAcylation in supporting longevity^64^ may render longer-lived immune cells such as T cells and resident macrophages more susceptible to loss of O-GlcNAcylation, compared to more rapidly replaced DC populations. Functionally, we found a selective impairment of DCs to prime Th2 responses following loss of OGT in different models of type 2 immunity, with consequences for disease outcome. This seems to be selective for Th2 responses as type 1 immunity or regulatory T cell responses were not affected in the models we tested. Finally, we extended our observations to primary DCs in humans by showing that cDC2s from individuals taking part in a controlled human hookworm infection trial displayed increased intracellular O-GlcNAcylation levels during the peak of their type 2 immune response. A change in O-GlcNAcylation was only consistently observed in CD1c^+^ cDC2s, but not cDC1s or pDCs. As CD1c^+^ DCs are the primary subset linked to CD4^+^ activation and polarization in humans^65, 66, 67^, our data provides a first indication that O-GlcNAcylation may also be important for DCs to drive Th2 responses in the context of human helminth infection.

A limitation of our current work is that we have not identified the specific O-GlcNAcylation sites on Zyxin and Fascin-1 through which the effects are mediated. Our MS platform did not allow us to reliably predict this. Further studies are needed to map these sites and assess their functional contribution to the phenotype. In addition, we found loss of O-GlcNAcylation in DCs resulted in a specific reduction in frequencies of these cells in the lung. Therefore we cannot exclude the possibility that specifically in the lung also limited antigen presentation contributes to the reduced HDM-induced type 2 immune response we observed in these mice, rather than only a DC-intrinsic defect in Th2 priming. Finally, it is possible that some of the observed metabolic reprogramming and dependencies of moDCs conditioned for Th2 priming *in vitro* do not fully translate to those of primed cDC2s in vivo. Hence, further studies are warranted for more in depth temporal and spatial exploration of how glycolysis shapes DC function during Th2 priming *in situ*.

In summary, we identified O-GlcNAcylation and low glycolysis as key metabolic checkpoints that uniquely define Th2-priming DCs and underpin their ability to prime this response. As both processes are highly sensitive to nutrient availability, it will be highly valuable in future studies to explore the interplay between local changes in tissue nutrient composition during type 2 infections and/or under other pathological settings, such as obesity, and these metabolic processes in DCs and how that impacts their Th2-priming capacity. This may shed new mechanistic light on the described positive association between obesity and allergies^68^, as well as increased resistance to helminth infections^69^. In addition, these unique metabolic characteristics of, and requirements for Th2 priming by DCs, provide an opportunity for selective manipulation of type 2 immune responses that can be harnessed to treat type 2-mediated inflammatory diseases.

## Methods

### Mice

*Itgax*^cre^ mice were crossed to *Ogt*^flox^ mice and bred in-house under specific pathogen free (SPF) conditions. For *in vivo* experiments only males were used, between an age of 8 and 16 weeks. Experiments were performed in accordance with local government regulations, EU Directive 2010/63EU and Recommendation 2007/526/EC regarding the protection of animals used for experimental and other scientific purposes, as well as approved by the Dutch Central Authority for Scientific Procedures on Animals (CCD). Animal license number AVD1160020198846.

### *Heligmosomoides polygyrus* infection

Mice were infected with *H. polygyrus* with 200 L3 larvae by oral gavage. Tissues were harvested 14 days post infection for immunological analysis. Fecal egg count was performed on feces of 21 days infected mice, and it was calculated as egg per gram (epg) of the fecal material. In brief, feces were dissolved in 2 mL water and left in the fridge overnight. The next day, 2 mL of saturated salt solution was added and mixed thoroughly with pasteur pipette. Approximately 150 uL of suspension was introduced in a McMaster chamber and viewed under a light microscope (10 x objective). The epg was calculated according to the equation: number of eggs counted/volume counted/weight of fecal material*total volume.

### *Schistosoma mansoni* infection

Mice were infected with *S. mansoni* (Puerto Rican strain; Naval Medical Research Institute) by 30 minutes of percutaneous exposure to 35 cercariae on shaved abdomen. To this end, mice were anesthetized with 50 mg/kg bodyweight ketamine + 10 mg/kg bodyweight xylazine. Female mice were assisted in waking up by i.p. injection with 150 µL of 0.4 mg/kg bodyweight. All injections were done using PBS and a 25G needle. All anesthetics were purchased at the LUMC pharmacy. Mice were culled 8 weeks later. Livers and hepatic LNs were collected to egg counts and immune profiling.

### House dust mite-induced allergic asthma model

Allergic airway inflammation was induced by sensitizing mice via intranasal administration of 1 μg HDM (Greer, London, United Kingdom) in 50 μL of PBS. One week later, these mice were challenged for 5 consecutive days via intranasal administration of 10 μg HDM in 50 μL of PBS. On day 15, bronchoalveolar lavage (BAL) fluid and lung draining mediastinal LNs (med LNs) were obtained to determine inflammatory cell recruitment. BAL was performed by instilling the lungs with 3 × 1mL aliquots of sterile PBS (Braun, Oss, The Netherlands. Eosinophilia was assessed in BAL by flow cytometry as a readout for allergic inflammation.

### Immunizations

For evaluation of T cell priming, mice were injected s.c. either with 25 μg of CpG-B with E7-SLP (a synthetic 43-mer-long peptide from the human papillomavirus (HPV) E7 protein that contains the immunodominant and H-2Db-restricted epitope RAHYNIVTF) or 10ug SEA (*S. mansoni* soluble egg antigen; for description of preparation see below) in the hind footpad. Mice were culled 7 days later and draining popLNs were collected for evaluation of T cell responses. To assess O-GlcNAcylation status of Ag^+^ DCs following immunizations with SEA, SEA was fluorescently labelled with Dylight® 488 Conjugation Kit (Fast) – Lightning-Link® according to the manufacturers protocol, after which 1ug was subcutaneously injected into the ear pinnae. Mice were culled 48h later and draining LNs were collected for evaluation of DC O-GlcNAcylation status. Contralateral LN served as negative controls.

### FITC painting

Mice were painted with 20 μL of inflammatory FITC paint (5 mg/mL fluorescein isothiocyanate [#F3651, Sigma] in a 1:1 mix of dibutylphalate [#524980, Sigma] and acetone [#100014, Merck, Amsterdam, The Netherlands]) on shaved flanks and draining ingLNs were collected either 2 days later.

### Organ processing

#### Intestines

The first 15cm of the small intestine was excised, opened longitudinally over PBS-soaked paper towel. Feces and mucus were gently scraped off with a metal spatula then intestine were washed by vigorous shaking in a 50 ml tube containing 15 ml of Ca/Mg-free HBSS and 2mM EDTA, then cut into 1-2cm pieces and stored in wash buffer on ice until further processing. Three rounds of washing were done in 15ml HBSS/EDTA wash buffer for 20min shaking at 200rpm for 20min. Washed intestines were rinsed once in RPMI before digestion for 15-20 minutes in 10 ml RPMI containing 10% FBS, 1 mg/ml collagenase VIII (Sigma, C2139) and 40U/ml DNAse I. Digestion was stopped by adding cold RPMI containing 10% FBS, then resulting suspension was filtered through a 100µM filter, spun at 400g for 5 min, resuspended in PBS/2%FBS/2mM EDTA and filtered through a 40µM filter for counting. Large intestines, including caecum, were processed similarly but digested with 1 mg/ml collagenase IV, 0.5 mg/ml collagenase D, 1 mg/ml dispase II, 40U/ml DNAse I for 25–30 min.

#### Livers

Livers were stored in cold RPMI then minced using a razor blade in a petri plate, and digested similarly to the large intestine with additional manual shaking vigorously by hand every 5–8 min. Digested livers were filtered through 100um strainers, and washed twice with PBS/2%FCS/2mM EDTA, and red-blood cell lysed with ACK lysis buffer. CD45+ cells were enriched by positive magnetic selection.

#### Lungs, Spleens, and Lymph Nodes

Lungs and spleens were stored in cold PBS, minced with scissors and digested in 1 mg/ml collagenase IV and 40U/ml DNAse I and shaken for 30 min before mashing through a 100 μm filter. Homogenates were red-blood cell lysed, washed, and passed through a 40μm. Lymph nodes were digested in 0.5mg/ml collagenase IV and 40U/ml DNAse I and otherwise processed similarly but without red blood cell lysis.

### Analysis of murine T cell responses

For detection of T cell cytokines, total cell homogenates were stimulated with PMA and Ionomycin in the presence of brefeldin A for 4h at 37 degrees. Stimulated cells were fixed with 1.8% formaldehyde in PBS for 15 minutes at RT and permeabilized with 1x eBio perm buffer. Staining for surface markers and intracellular cytokines was performed in 1x perm buffer. Stained cells were washed twice and resuspended in FACS buffer for acquisition.

### Preparation of *Schistosoma mansoni* soluble egg antigens and omega-1

*S. mansoni* eggs were isolated and processed into a SEA preparation as described previously^28^. Protein concentration was determined using a bicinchoninic acid (BCA) protein assay kit (Pierce, #PIER23225). Endotoxin contamination was determined by a direct comparison of SEA batches to LPS in a TLR4-transfected Human Embryonic Kidney 293 (HEK) reporter cell line, Omega-1 was purified from SEA as described^28^ or produced and glyco-engineered in plants as described^70^.

### Isolation of peripheral blood mononuclear cells

Peripheral venous blood, from buffycoats provided by Sanquin (Amsterdam, The Netherlands), was divided in conical 50 mL tubes (#227261, Greiner) with 20 mL of room temperature venous blood diluted 1:1 with room temperature HBSS (#14170-088, Gibco) per tube. This solution was gently mixed before addition of 13 mL room temperature Ficoll (#17-1440-02, GE healthcare) underneath the blood:HBSS. Alternatively, all liquids were used cold, but temperatures were never mixed. Peripheral blood mononuclear cells (PMBCs) were concentrated into a ring by density centrifugation at 400g with low brake for 25 minutes at room temperature. The rings were transferred using a Pasteur pipette (#861172001, #Sarstedt) to new 50 mL tubes to which HBSS supplemented with 1% v/v heat-inactivated foetal calf serum (HI-FCS; #S-FBS-EU-015, Serana, Pessin, Germany) was added up to 50 mL. Blood platelets were removed by two rounds of centrifugation at 200g with normal brake for 20 minutes at room temperature. Finally, the PBMCs were resuspended in PBS supplemented with 0.5% BSA [fraction V, #10735086001, Roche, Woerden, The Netherlands] and 2 mM EDTA [#15575-038, Thermo] that was sterilized using a 0.22 μm filter system (#431097, Corning).

### Monocyte-derived dendritic cell culture

PBMCs were counted using an inhouse Türk solution (= 50 mg gentian violet, 5 mL glacial acetic acid and 500 mL H20) and brought to a concentration of 10^7 cells per 95 µL of MACS buffer. Monocytes were isolated by addition of 5 µL of CD14 MACS beads (#120-007-943, Miltenyi) per 10^7 cells, mixing well, incubating for 15 minutes at 4 degrees Celsius and column sorting using LS columns (#130-042-401, Miltenyi) according to the manufacturer’s protocol provided with the beads. Whole PBMCs, CD14- flow through and CD14^+^ monocyte fractions were set aside to check for purity, which routinely was >95%. Monocytes were brought to 400 cells per µL in ice-cold monocyte-derived dendritic cell (moDC) differentiation medium (= no additives RPMI [naRPMI; #42401-042, Invitrogen] supplemented with 10% HI-FCS, 2 mM L-glutamine [#G-8540-100g, Sigma], 100 U/mL penicillin [#16128286, Eureco-pharma, Ridderkerk, The Netherlands] and 100 µg/mL streptomycin [#S9137, Sigma] that together formed our general human immune cell RPMI for metabolism work and also 10 ng/mL of human recombinant granulocyte/macrophage colony-stimulating factor (GM-CSF; #PHC2013, Biosource) and 0.86 ng/mL of human recombinant IL-4 (#204-IL, R&D systems) was added for moDC differentiation. The media was kept cold to prevent monocytes sticking to the plastic tube. Monocytes were usually plated at 2 million cells per 5 mL of media in 6-well plates (#140675, Nunc) and after 2 or 3 days, the top half of the medium was carefully removed and replaced by 2.5 mL of fresh medium complemented with 20 ng/mL of GM-CSF and 1.72 ng/mL of IL-4 to maintain the concentrations of differentiating cytokines.

### Lactate assay

Lactate concentration in supernatants was determined by measuring absorbance of NADH at 340 nm after the reaction of lactate with NAD by lactate dehydrogenase into pyruvate and NADH. To this end, 10ul sample was mixed with reaction mix (NAD 0.75 mM [Roche Applied Science 10127965001], Glycine 0.4 mM [SIGMA 410225], Hydrazine hydrate 0.4 M [SIGMA 225819] and L-Lactate Dehydrogenase from rabbit muscle [Roche Applied Science 10127876001] in milliQ H_2_O. Sodium L-lactate [SIGMA L7022] was used a as standard.

### siRNA-based gene silencing

At day 4 of the culture, the cells were harvested, washed with PBS, brought to a concentration of 1.1x10^6 cells per 100 µL and split into 1.5 mL Eppendorf tubes with 100 µL of cell solution. Shortly before transfection by electroporation, tubes were spun down, cell pellets were carefully pipetted dry, pellets were reconstituted in 100 µL of resuspension buffer and 10 µL of 455 nM siRNA was added (On-Targetplus Smartpool for OGT [#M-019111-00-0005], ZYX [#L-016734-00-0005], FSCN1 [#L-019576-00-0005], or scrambled control [#D-001206-13-05, Darmacon, Lafayette, Colorado, United States Darmacon]). Electroporation as described here was performed using the 100 µL variant (#MPK10096, Invitrogen) of the Neon Transfection System (#MPK5000, Invitrogen) and a complementary pipette (#MPP100, Invitrogen) with the following settings: 1600 V, 20 ms and one pulse. Immediately after electroporation, the cells were transferred to 5 mL of 10% HI-FCS RPMI-1640 with 2 mM glutamine but without further additions. Importantly, the media contained no antibiotics at this stage. Electroporation tips were re-used to a maximum of three times for the same target. Later, the cells were plated at 200 cells per µL in, if cell numbers allowed it, 6-well plates again. The next morning, the media was re-supplemented with penicillin, streptomycin, rGM-CSF and rIL-4. At day 6, the cells were handled as normally. Reconstituting the pellet in a total volume of 110 µL helped prevent sucking up air bubbles into the electroporation tip, which is crucial. Minimizing time spend in the resuspension buffer helped with cell viability. Silencing efficiency was determined by qPCR on 6 days old cells and was routinely greater than 80%.

### DC stimulation

Day 5-7 moDCs were harvested without discarding the differentiation media and replated in 96-well plates (#167008, Nunc) at 5-10x10^4 cells per 100 µL of the same media and left to rest for 2-3 hours. Alternatively, cells were plated in 200 µL and the next day the top half of the medium was carefully removed. Inhibitors were added 30 minutes prior to antigen stimulation and the cells were kept for a total of 24 hours in a 5% CO2 and 37 degrees Celsius cell incubator. Inhibitors included: 5-10 mM of 2-deoxyglucose (2-DG; in mQ water; #D8375-1g, Sigma), 20 µM ST045849 (ST; in DMSO; #6775, Tocris) and 10 µM Thiamet G (TG; in DMSO; #13237, Cayman Chemical, Ann Arbor, Michigan, United States). Antigens included 100 ng/mL of ultrapure lipopolysaccharides (LPS; *Escherichia coli* 0111 B4 strain, #tlrl-3eblps, InvivoGen), 20 µg/mL of polyinosinic:polycytidylic acid (PolyIC; high molecular weight, #TLRL-PIC, InvivoGen), 20-50 µg/mL of zymosan (#Z4250, Sigma), 250-500 ng/mL of natural omega-1, 2 µg mL of plant-derived recombinant omega-1, 25-50 µg/mL of soluble egg antigen (SEA); and 10 µg/mL of house dust mite (HDM; Greer, lot number: 305470). Supernatants were collected after 24 hours and IL-12p70 concentrations were determined using ELISA (BD #555183). Alternatively, 10.000 24h stimulated DCs were co-cultured with 10.000 J558-CD40L cells - a CD40L expressing cell line – for another 24 hours before determination of cytokine concentrations. Viability, differentiation makers and expression of costimulatory molecule expression was determined using flow cytometry on a FACSCanto II.

For RNA sequencing and metabolomics analyses, moDCs from three 6-well plates were pooled, centrifuged and resuspended in 1 mL himRPMI. 300 µL was used for mRNA extraction, 300 µL was used for metabolite extraction, 300 µL was used for extracellular flux (XF) analysis and the remaining 100 µL was used for cell counting, analysis of DC activation and T cell polarization. Cell numbers for mRNA extraction and metabolite extraction ranged between 0.3-1.1 million cells. mRNA was extracted with oligo-dT beads (Invitrogen) and libraries were prepared and quantified as described before^71^. Extraction of metabolites was done according to the recommendations of General Metabolics (Boston, Massachusetts, USA) specifically for extraction of polar metabolites from adherent mammalian cell culture. Briefly, cells were transferred to 1.5 mL Eppendorfs, centrifuged, washed using a warm ammonium carbonate wash solution (75 mM ammonium carbonate [#A9516, Sigma] in HLPC-grade water and pH 7.4 titrated with HLPC-grade formic acid) and resuspended in a 70 degrees Celsius 70% ethanol extraction solvent (70% v/v absolute ethanol [#100983.1000, Merck] in HLPC-grade water). After exactly 3 minutes, the extracts were transferred to other, pre-chilled Eppendorfs in a dry-ice bed. The first group of Eppendorfs were washed with a second volume of extraction solvent that was immediately transferred to the second group of Eppendorfs. Combined extracts were then centrifuged at 14.000 rpm for 10 minutes at 4 degrees Celsius or colder, after which a fixed volume of the extract supernatant was collected – without disturbing any pelleted material – and transferred to a third group of Eppendorfs, also pre-chilled and in a dry-ice bed. The materials were stored at -80 degrees Celsius before shipping processing by General Metabolics as previously described^27^.

### *In vitro* T cell polarization

Naïve CD4 T cells for assessment of DC capacity for Th1/2 polarization were isolated from allogenic buffy coat PBMCs using a naïve human CD4 T cell isolation kit (#480042, BioLegend). Memory CD4 T cells for assessment of DC capacity for Th17 reactivation were isolated using a human CD4 T cell isolation kit (#130-096-533, Miltenyi) followed by a combination of anti-CD45RO-PE (UCHL1 clone; #R0843, Dako) and anti-PE microbreads (#130-048-801) to eliminate naïve CD4 T cells. For Th1/2 polarization, 5.000 DCs that were stimulated for 24h were co-cultured with 20.000 allogenic naïve CD4 T cells in the presence of 20 pg/mL Staphylococcal Enterotoxin B (SEB; #S4881, Sigma) in a cell culture treated flat-bottom 96 well plate. After 5-7 days, cells were transferred to a flat-bottom 24 well plate well containing 1 mL of media that was supplemented with 42 IU/mL recombinant human IL-2 (#202-IL, R&D systems). 2 days later, 1 mL of fresh media with IL-2 was added and cell cultures were split into two. Cells were restimulated at approximately 11 days with 100 ng/mL phorbol myristate acetate (PMA) and 2 µg/mL ionomycin for a total 6 hours, of which the last 4 hours in the presence of 10 µg/mL brefeldin A (#B-7651, Sigma). Cells were stained with Aqua LIVE/DEAD and fixed with formalin as described in the next section before permeabilization with a permeabilization buffer (#00-8333-56, Thermo) according to the manufacturer’s recommendation. Production of IL-4 and IFNγ was determined using flow cytometry. For Th17 reactivation, 2.000 stimulated DCs were co-cultured with 20.000 allogenic memory CD4 T cells in IMDM (#12-722F, Lonza) supplemented with 10% HI-FCS, 100 U/mL penicillin and 100 µg/mL streptomycin in the presence of 10 pg/mL SEB in a flat-bottom 96 well plate. Supernatants were collected after 5 days and IL-17 concentration was determined using ELISA.

### Analysis of blood DCs from hookworm controlled human infection trial

Peripheral venous blood was drawn from healthy volunteers prior to and 8 weeks after infection with *Necator americanus*, as part of a controlled human infection trial (NCT03126552)^42^. PBMCs were isolated as described above and cryopreserved in liquid nitrogen until later analysis.

### Flow cytometry

#### Human cells

A complete list of antibodies and dilutions is shown in Supplemental **Table 3**. In general, human moDC single cell suspensions underwent viability staining for 20 minutes at room temperature using the LIVE/DEAD^TM^ Fixable Aqua Dead Cell Stain Kit (#L34957, Thermo) 1:400 in PBS and fixation for 15 minutes at room temperature using 1.85% formaldehyde (F1635, Sigma) in PBS before surface staining with antibodies in FACS buffer (PBS, 0.5% BSA, 2mM EDTA) for 30 minutes at 4 degrees Celsius. Alternatively, cells were acquired unfixed in FACS buffer with 7-AAD (#00-6993-50, eBioscience) 1:50 to distinguish live from dead cells. Compensation was done using single staining on beads (#552843, BD Biosciences or 552845, BD Biosciences or #A10346, Invitrogen). For detection of protein O-GlcNAcylation in moDCs, cells underwent Aqua viability staining for 30 minutes on ice and fixation for 60 minutes at 4 degrees using the Foxp3 / Transcription Factor Staining Buffer Set (#00-5523-00, eBioscience). Afterwards, cells were further permeabilized by first resuspending them in 20 µL of ice-cold PBS and then adding 180 µL of ice-cold absolute methanol (90% end concentration; #1.06009, Merck). Methanol was added carefully to prevent leaking of methanol out of the pipette tips and overflowing of the well plates. Cells were fixed for at least 10 minutes at -20 degrees Celsius up to overnight. Methanol was washed away with FACS buffer at least two times before staining O-GlcNAcylation, CD1a and CD14 to prevent interaction with methanol sensitive fluorochromes.. To assess levels of O-GlcNAcylation in DCs subsets in frozen human blood, PBMCs were thawed and first live stained for the following surface markers during the fixable Aqua stain: CD3/CD56/CD19, which were used as exclusion markers, and HLA-DR and CD11c, which in combination with CD123, CD141, and CD1c were used to identify pDCs, cDC1s and cDC2s respectively. Cells were then washed in PBS and fixed with an Intracellular Fixation & Permeabilization Buffer Set (#88-8824-00, eBioscience) for 1 h at 4 degrees Celsius. Cells we washed again in PBS and subsequently permeabilized using the same set before a second staining for O-GlcNAcylation as described above. Cells were analyzed on an LSR-II.

#### Murine cells

Murine cells were plated in a 96-well V-bottom plate, washed with PBS and pre-stained with viability dye and Fc-block in PBS for 15 min at RT. If required, subsequent live surface-staining was performed for fixation sensitive targets in FACS buffer for 30 min on ice before fixation. Fixation was performed using eBioscience Foxp3 fixation/permeabilization staining kit for 30-60min at RT. Fixed cells were stored in FACS buffer at 4 degrees until further staining, at which point cells were washed with 1x permeabilization buffer and resuspended in staining mix for 1h at RT. Staining mixed contained the remaining antibodies diluted in perm buffer 1x Brilliant Stain Buffer Plus (BD Biosciences, 566385) and TrueStain Monocyte Block (BioLegend, 426103). Stained cells were washed twice with perm buffer and resuspended in FACS buffer for acquisition. For the preservation of Ag+ DCs after subcutaneous immunisation of fluorescent antigens, fixation was performed in methanol containing PFA 1% for 20min on ice. Fixed cells were then incubated overnight at 4 degrees with antibodies diluted in FACS buffer containing 0.025% saponin for intracellular detection of selected targets. Samples were acquired on a FACSCanto II, LSR-II (both BD Biosciences) or Aurora 5-laser spectral flow cytometer (Cytek). Acquired samples were compensated or unmixed using using FlowJo (Version 10, TreeStar, Meerhout, Belgium) and SpectroFlo version 3, and analyzed with FlowJo and OMIQ.

### O-GlcNAcylome analysis

#### Preparation of proteins

6h after stimulation 2 million human moDCs were lysed in 400 µL of RIPA lysis buffer (25 mM Tris HCl, 150 mM NaCl, 1% NP-40, 1% sodium deoxycholate, 0.1% SDS at pH 7.6) supplemented with a protease/phosphatase inhibitor cocktail (ThermoFisher) and 1 µM of O-GlcNAc cycling enzyme inhibitors (Sigma-Aldrich, Saint-Louis, Missouri, United-States). Protein contents were determined using BCA kit (add reference + manufacturer) and 250 µg of proteins from the lysate were then precipitated using chloroform/methanol (MeOH) precipitation method as follows. Increase volumes accordingly if the volume of the input material exceeded 200 uL. 600 μL of MeOH were added to the 200 μL sample, followed by 150 μL of chloroform and 400 μL of 18 megaOhm water. Vortex between each addition.Centrifuge for 5 min at 13,000×g. The upper aqueous phase was carefully removed and discarded. Additional 450 μL MeOH were added to pellet the protein after brief vortex and centrifugation for 5 min at 13,000×g 4◦C. The supernatant was then removed, and the pellet air dried for 5 min. Finally, proteins were resuspended in 40 µL of 1% SDS in 20 mM HEPES pH 7.9 and 1% SDS pH 7.9 and heated 5-10 min at 90 °C to assure completely resuspension of proteins. Finally, proteins were shortly cooled on ice.

#### Enzymatic labelling and O-GlcNAcylated protein enrichment

O-GlcNAc groups from proteins were labelled with tetramethylrhodamine azide (TAMRA) using the Click-iT® O-GlcNAc enzymatic labelling system kit (C33368) followed by the Click-iT® protein analysis detection kit (C33370) from Invitrogen according to the manufacturer’s instructions. Next, lysates were subjected to another chloroform/MeOH precipitation as described previously. Samples were resuspended at 2 mg/mL in Tris-HCl (50 mM) 1% SDS, pH 8.0. After heating proteins for resuspension, SDS was quenched with similar volume of NEFTD buffer (100 mM NaCl, 50 mMTris-HCl, 5 mM EDTA, 6% NP-40 at pH 7.4). Before immunoprecipitation of TAMRA labelled proteins, lysate was precleared with washed protein G sepharose beads (Sigma-Aldrich, GE17-0618-01) in PBS for 1h at 4◦C shaking vigorously to avoid non-specific binding of proteins on the beads. Afterwards, supernatant was incubated with pre-washed protein G sepharose beads (10 µL/500 ug of protein) coupled with anti-TAMRA antibody (15 µg/10uL beads, A6397, Invitrogen) for 1.5 h at 4 °C. shaking vigorously Following centrifugation (500×g, 1 min), the beads were washed once with NEFTD buffer and three times with NEFT buffer (NEFTD without NP-40). The beads were then boiled 5 min in Laemmli buffer (2 mM EDTA, 4% SDS, 20% Glycerol, 0.004% bromophenol blue, 50 mM DTT, and 100 mM Tris at pH 6.8) to elute O-GlcNAc proteins. Proteins were retrieved after centrifugation (13000g, 5 min, RT).

#### In-Gel Digestion and identification of captured O-GlcNAcylated Proteins

For MS analysis, gel bands were reduced with 10 mM dithiothreitol, alkylated with 50 mM iodoacetamide. In-gel trypsin digestion was performed using a Proteineer DP digestion robot (Bruker). Tryptic peptides were extracted from the gel slices using 50/50/0.1 water/acetonitrile/formic acid (v/v/v), after which samples were lyophilized. The samples were dissolved in water/formic acid (100/0.1 v/v) and analyzed by on-line C18 nanoHPLC MS/MS with a system consisting of an Ultimate3000nano gradient HPLC system (Thermo, Bremen, Germany), and an Exploris480 mass spectrometer (Thermo). Peptides were injected onto a cartridge precolumn (300 μm × 5 mm, C18 PepMap, 5 um, 100 A, and eluted via a homemade analytical nano-HPLC column (30 cm × 75 μm; Reprosil-Pur C18-AQ 1.9 um, 120 A (Dr. Maisch, Ammerbuch, Germany). The gradient was run from 2% to 40% solvent B (20/80/0.1 water/acetonitrile/formic acid (FA) v/v) in 30 min. The nano-HPLC column was drawn to a tip of ∼10 μm and acted as the electrospray needle of the MS source. The mass spectrometer was operated in data-dependent Top 20 MS/MS mode, with a HCD collision energy at 30% and recording of the MS2 spectrum in the orbitrap, with a quadrupole isolation width of 1.2 Da. In the master scan (MS1) the resolution was 120,000, the scan range 400-1500, at standard AGC target @maximum fill time of 50 ms. A lock mass correction on the background ion m/z=445.12003 was used. Precursors were dynamically excluded after n=1 with an exclusion duration of 10 s, and with a precursor range of 20 ppm. Charge states 2-5 were included. For MS2 the first mass was set to 110 Da, and the MS2 scan resolution was 30,000 at an AGC target of 100% @maximum fill time of 60 ms. In a post-analysis process, Proteome Discoverer 2.5 (Thermo) was used with 10 ppm and 20 ppm deviation for precursor and fragment mass, respectively and trypsin as enzyme was specified. Oxidation on Met and acetylation on N-term were set as a common modification; carbamidomethyl on Cys was set as a fixed modification. An FDR of 1% was set. The mass spectrometry proteomics data have been deposited to the ProteomeXchange Consortium via the PRIDE partner repository with the dataset identifier PXD073285.

#### Pathway analysis and bioinformatics

The analysis was conducted in RStudio. Missing abundance values were set at 50.000, half of the noise cutoff. The differential expression between control and stimulated DCs was statistically assessed through a linear mixed model using the lmer function from the lmerTest package, with log10-transformed abundance as the outcome variable, stimulation and stimulation time as the predictor variables, and random intercepts for the donors. Proteins with p-values lower than 0.05 were considered for protein-protein interaction and pathway enrichment analysis. Protein-protein interaction scores from the STRING database were retrieved using the get_interactions function from the STRINGdb package. Database version was 12.0, minimum score threshold was 400 and both physical and functional interactions were included. Interactions were plotted as an undirected network with a Fruchterman-Reingold force-directed layout. Reactome database pathway enrichment was performed using the get_enrichment function from the STRINGdb package, which computes enrichment using an hypergeometric test and calculates false discovery rate using the Benjamini-Hochberg procedure. Enriched pathways were expressed as Rich Factors, which is the ratio of mapped proteins tot total proteins in a pathway. All plotes were made using the ggplot function from the ggplot2 package.

### Confocal microscopy

On day 6 of human moDC differentiation, cells were seeded in glass bottom confocal dishes (627871; Greiner) and coated for 2 h with poly-D-lysine (PDL) at 37°C. Treatments were performed as described earlier and DCs were analyzed 24h later. Cells were washed 3x with 300 ul PBS. Next cells were fixed with 300 ul 2% PFA for 15 min at RT. After 3 washes in PBS, cells were permeabilized for 15 min with 300 uL PBS containing 0.1% Triton X-100. After 3 washes with 300 ul PBS with 0.01% Tween, cells were treated with 300 ul 1% BSA/PBST (PBS+ 0.1% Tween 20) for 30 min to block unspecific binding of antibodies. Cells were washed 3x with 300 ul PBST and subsequently incubated with 100 uL antibody stain (1:100 Zyxin-biotin (ab109316; Abcam), 1;400 Phalloidin-FITC (ab235137; Abcam) and 1:200 Hoechst [33342 (H3570; Thermo Fisher Scientific)) in 1% BSA/PBST for 30 min at RT in the dark, followed by 1:200 streptavidin-BV605 staining (405229; Biolegend) in 1% BSA/PBST. After Washes 2x with 300 ul PBST and once with PBS cells were stored in the dark until imaging. Cells were imaged directly using the 40× objective on a SP8X WLL (white light laser) microscope (Leica Microsystems). Images were taken with focal plane close to site of adherence to visualize podosomes. Single cell images were made by using Photoshop for both fluorescent signals (Zyxin=red, Xyxin=green). For each cell the different colored images were merged in ImageJ-Fiji and the colocalization of both signals were analyzed with the BIOP JACoP plugin. Standard settings were used with setting a default threshold, that was based on the unstimulated sample of each donor.

### Atomic Force Microscopy (AFM)-based single-cell force spectroscopy (SCFS)

The experiments were performed as previously described using a JPK CellHesion unit^41^. In brief, To measure T–DC adhesion forces, DC2.4 transduced with CIITA to increase MHCII expression, were cultured on glass disks for at least 24 h before experiments.. In certain experiments, DC2.4 cells were pulsed with 10 µg/mL soluble OVA protein and treated with 100 ng/mL LPS or Omega-1, then cultured overnight. The OSMI group received treatment 1 h ahead of the other groups. OTII mouse CD4^+^ CD25^-^T cells were isolated with CD4 and CD25 magnetic beads, and then co-cultured overnight with CIITA-overexpressing DC2.4 cells in the presence of 10 μg/ml OVA and 200 IU/ml IL-2.The disks were moved into an AFM-compatible chamber and mounted on to the machine stage. A clean cantilever was coated with Cell-Tak (BD), and then used to glue individual OTII Th cells added to the disk. In each cycle, the AFM cantilever carrying a single OTII Th cell was lowered to allow T cell contact with an individual DC by 0.5–2-µm increments until the first force curve was generated. The T cell on the cantilever was then allowed to interact with the DC for 15 s before being moved upwards, until two cells were separated completely. The process was then repeated. The incubator chamber in which the machine was housed was conditioned at 37°C and at 5% CO_2_. In all experiments, a minimum of 14 force curves were collected for further analysis. For each SCFS experiment, a pair of T–DC was used to generate force readings from each up and down cycle over a period of several minutes. At least three such pairs were used for each condition. All data from these readings (at least three pairs) were used as a group for comparison among the groups. This analysis is considered as one experiment For selected experiments, *Fascin-1 (Fscn1)* and *Zyxin (Zyx)* were knocked out in DC2.4 cells using a CRISPR/Cas9-based approach. Single guide RNAs (gRNAs) targeting *Fscn1* or *Zyx* were designed and cloned into the lentiCRISPR v2 vector. Briefly, lentiCRISPR v2 was digested with Esp3I, and the ∼11 kb backbone fragment was gel-purified. Complementary gRNA oligonucleotides were phosphorylated, annealed, and ligated into the digested vector. Ligation products were transformed into competent bacteria, and positive clones were verified by Sanger sequencing using the hU6-F primer. Correct plasmids were purified for subsequent lentivirus production. The gRNA sequences used were as follows: *Fscn1*: Oligo 1: 5′-CACCGTGTCCAGTACTTGCCCGTGT-3′, Oligo 2: 5′-CACAGGTCATGAACGGGCACACAAA-3′. *Zyx*: Oligo 1: 5′-CACCGCTGCGGGGCGTAAAACGCGG-3′, Oligo 2: 5′-CGACGCCCCGCATTTTGCGCCCAAA-3′. Lentiviral particles were produced by co-transfecting L293 cells with the CRISPR plasmid, psPAX2, and pMD2.G using Neurofect transfection reagent, according to the manufacturer’s instructions. Viral supernatants were collected, concentrated, and used to infect CIITA-overexpressing DC2.4 cells in the presence of polybrene. After 14–16 h, the medium was replaced with fresh complete medium. To establish stable knockout lines, the minimal lethal concentration of puromycin was first determined in wild-type DC2.4 cells (2.5 µg/mL). Infected cells were then selected with puromycin at this concentration. Surviving cells were harvested, diluted, and seeded at approximately 5 cells/mL into 96-well plates to allow clonal expansion of single-cell–derived colonies. Knockout efficiency was assessed at both the genomic and protein levels. Genomic DNA was extracted from individual monoclonal populations, and target regions were PCR-amplified using gene-specific primers. PCR products were cloned into the pFAST-T vector and transformed into competent bacteria. Mutations were identified by Sanger sequencing using the M13-F primer. Loss of Fascin-1 or Zyxin protein expression was further confirmed by Western blot analysis.

The PCR primer sequences used for genomic validation were as follows: *Fscn1*: Forward: 5′-CCTGGAATCTGAGTGAGACA-3′, Reverse: 5′-GTAACTGGAACGGTTGGCA-3′; *Zyx*: Forward: 5′-CCGCCCCGGGAAGGGGACG-3′, Reverse: 5′-AGGAGGGGGCAAAGGAAAG-3′.

### RNA-seq and metabolic network analysis

RNA-seq libraries were sequenced using paired-end sequencing, second read (read-mate) was used for sample demultiplexing. Reads were aligned to the GRCh37 assembly of human genome using STAR aligner. Aligned reads were quantified using quant3p script (https://github.com/ctlab/quant3p) to account for specifics of 3’ sequencing with protein-coding subset of Gencode gene annotation. DESeq2 was used for differential expression analysis. Metabolic network clustering analysis was done with GAM-clustering method^72^ based on gene expression profiles and metabolic network based on KEGG database. Metabolic module for Th2-DC vs iDC comparison was done using Shiny GATOM method as previously described^73^, based on both DESeq2-based differential expression results for genes and limma-based differential analysis results for metabolites.

### Extracellular flux analysis

For real-time metabolic analysis on live cell, DCs were resuspended in Seahorse media (= unbuffered RPMI [#R1383-10X1L, Sigma] of which the pH was set to 7.4 using 37% HCl (#1.00317.1000, Merck) before sterilization using a 0.22 μm filter system that was supplemented with 5% HI-FCS and plated in XF cell culture plates (part of pack: # cat# 102416-100, Agilent, Amstelveen, The Netherlands) at 5x10^4 cells per 180 µL of Seahorse media. The plate was quickly spun down to facilitate the adherence of cells to the bottom of the well and then, the cells were rested for 1 hour in a special 0% CO2 cell incubator at 37 degrees Celsius. This rest time is also needed for the evaporation of remaining CO2 in the media, which can acidify the unbuffered media. To facilitate the adherence of semi-adherent DCs even more, the assay plate wells were coated beforehand with 25 µL of 50 µg/mL Poly-D-Lysine hydrobromide (PDL; diluted in mQ water; # A-003-E, Merck) for a minimum of 1 hour at 37 degrees Celsius, after which the PDL was pipetted away and the wells were washed using 200 µL of mQ water and subsequent air-drying. MilliQ water is preferred over PBS here as air-drying PBS residues might result in crystal formation. During the rest, pre-hydrated XF cartridges were filled with 10x concentrated assay reagents in Seahorse media that did not contain HI-FCS for sequential injections of glucose (10 mM end; port A; #G8644-100mL, Sigma), oligomycin (1 μM end; port B; #11342, Cayman Chemical), fluoro-carbonyl cyanide phenylhydrazone (FCCP, 3 μM end; port C; # C2920-10MG, Sigma), and finally, rotenone/antimycin A (1/1 μM end; port D; #R8875-1G/#A8674-25MG, Sigma). The manufacturer recommended the removal of HI-FCS from the injection solution to prevent clogging of the injection ports. Oxygen consumption rates (OCR) and extracellular acidification rates (ECAR) were recorded with the XF96e Extracellular Flux analyser (Agilent) and analyzed using Wave Desktop (Version 2.6, Agilent). Immediately after the run, XF culture plates were collected from the analyser, supernatants were carefully pipetted away, cells were washed gently with PBS, lysed in 25 µL of a RIPA-like buffer (50 mM Tris-HCl [pH 7.4], 150 mM NaCl and 0.5% SDS) and the plate was stored at -20 degrees Celsius for later protein content quantification using bicinchoninic acid protein assay kit (BCA; #PIER23225, Pierce) according to the manufacturer’s recommendations. Glycolytic rate = increase in ECAR in response to injection A. Glycolytic capacity = increase in ECAR in response to injection A and injection B. Baseline OCR = difference in OCR between readings following port A injection and readings after port D injection. Spare OCR is difference between basal and maximum OCR, which is calculated based on the difference in OCR between readings following port C injection and readings after port A injection. ATP production = difference in OCR between readings following port A injection and readings after port B injection. Proton leak = difference in OCR between readings following port B injection and after port D injection.

### Western blotting

DCs were washed twice with PBS before being lysed in EBSB buffer (8% [w/v] glycerol, 3% [w/v] SDS and 100 mM Tris–HCl [pH 6.8]). Lysates were immediately boiled for 5 min and their protein content was determined using a BCA kit. Protein was separated by SDS-PAGE followed by transfer to a PVDF membrane. Protein content of 1 million moDCs that were stimulated for 24h but not recounted before lysis in 150 µL was approximately 0.5 mg/mL. Usually 10 µg of protein was added to each lane. Membranes were blocked for 1 h at room temperature in TTBS buffer (20 mM Tris–HCl [pH 7.6], 137 mM NaCl, and 0.25% [v/v] Tween 20) containing 5% (w/v) fat free milk and incubated overnight with primary antibodies. The primary antibodies used were O-GlcNAc (RL2 clone; 1:1000, #MA1-072, Thermofisher), OGT (D1D8Q clone; 1:1000; #24083S, Cell Signaling and beta-actin (AC-15 clone; 1:10.000; #A5441, Sigma). The membranes were then washed in TTBS buffer and incubated with horseradish peroxidase-conjugated secondary antibodies for 2 h at room temperature. After washing, blots were developed using enhanced chemiluminescence.

### qPCR

RNA was extracted from snap-frozen pulsed DCs. The isolation of mRNA was performed according to manufacturer’s instruction using micro RNA Plus kit (Qiagen). cDNA was synthesized with reverse transcriptase kit (Promega) and PCR amplification by the SYBER Green method were done using CFX (Biorad). Specific primers for detected genes are as follows: human OGT forward is AGAAGGGCAGTGTTGCTGAAG and reverse is TGATATTGGCTAGGTTATTCAGAGAGTCT. The cycle threshold (Ct) value is defined as the number of PCR cycles in which the fluorescence signal exceeds the detection threshold value. The normalized amount of targeted mRNA (Nt) was calculated from Ct value obtained for both target and ACTB mRNA (served as housekeeping gene) with the equation Nt=2Ct(β-actin)-Ct(target). Relative mRNA expression was obtained by setting Nt in control as 1 in each experiment and for each donor.

### Quantification and statistical analysis

Statistical analysis as specified in figure legends were performed with Prism 9 (GraphPad software Inc.). Comparisons between two or more independent data groups were made by Student’s *t* test or analysis of variance test (ANOVA) with Fisher’s post-hoc test for multiple comparisons, respectively. Data distribution was assumed to be normal but this was not formally tested. Details of statistical tests are provided in the figure legends. p value < 0.05 was considered significant (* for p < 0.05, ** for p < 0.01 and *** for p < 0.001).

## Acknowledgements

This work was supported by an LUMC fellowship and VIDI grant (#91614087 by Netherlands Organisation for Scientific Research) awarded to BE. We thank the flow cytometry core at the LUMC for their support. We thank Gabriele Schram for providing natural omega-1.

## Author contributions

LP, MQ, JF, AH, FO, TP, GH, BW, JK, XW, JD, RT, AR and MF performed experiments. LP, MQ, JF, AS, MA, TP, GH and BE analyzed experiments. RM, GS provided reagents. LB, PV, CH, MA helped with experimental design and analysis. MR provided PBMCs from the controlled human hookworm infection trial. LP, MQ, TX, YS, OL and BE designed experiments. BE conceived and supervised the study and wrote the manuscript together with LP, JF and MQ.

## Data Availability

Data will either be made available or shared upon reasonable request following publication acceptance. Raw and processed transcriptomic data were deposited to Gene Expression Omnibus upon publication.

## Extended Data

**Extended Data Fig. 1:**
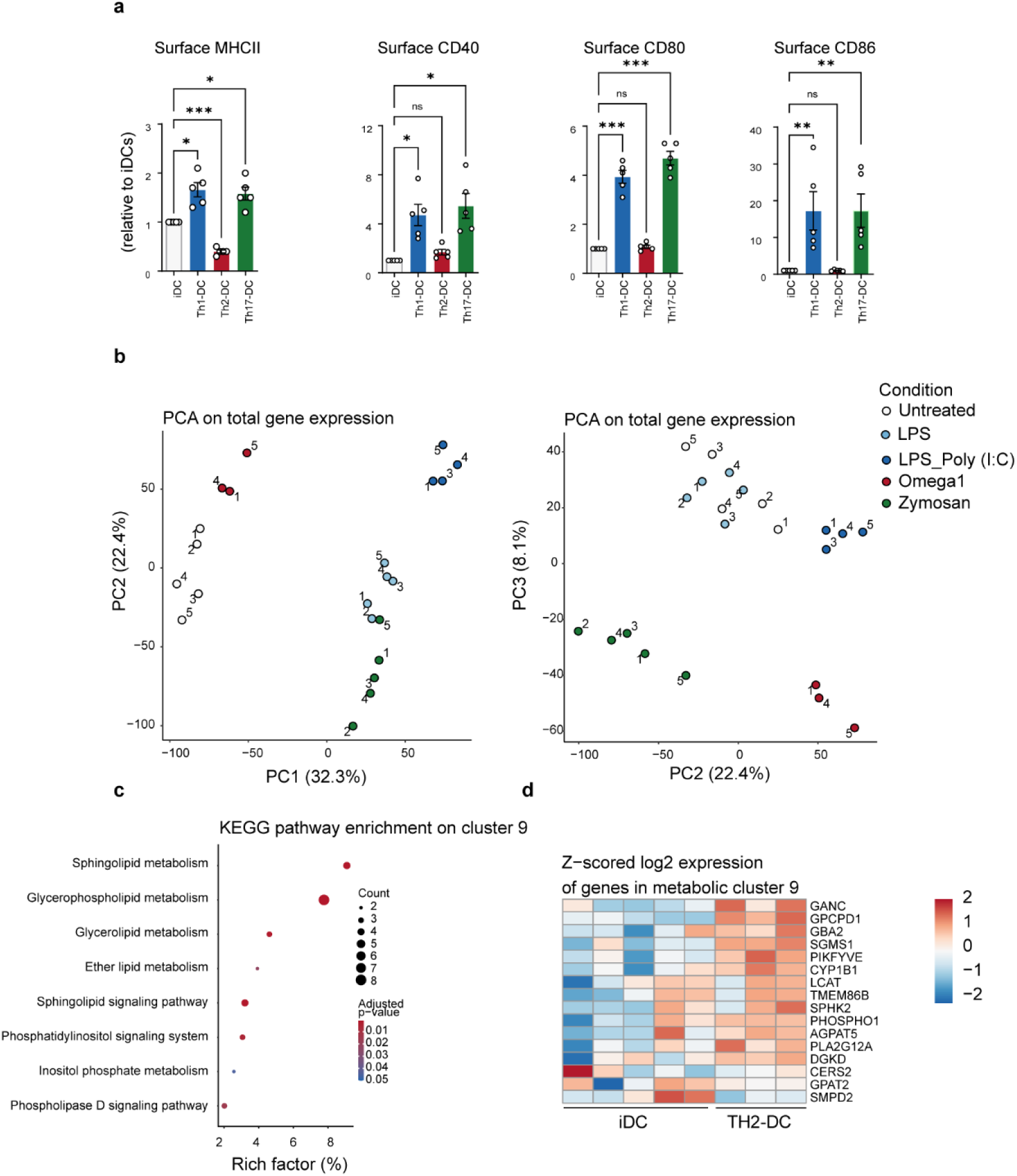
*In vitro* metabolic characterization of Th2-priming DCs. Human moDCs were left untreated (iDC in grey) or stimulated for 24h with either LPS+PolyIC (Th1-DC in light blue), omega-1 (Th2-DC in red) or zymosan (Th17-DC in green). **a**, The expression of maturation makers – based on the geometric mean fluorescence – are shown relative to iDCs. **b,** Principal component analysis on total transcriptome of differently conditioned DCs. The number of independent experiments is represented by symbols in the graphs and shown as mean ± SEM. **c,** KEGG pathways enrichment analysis of genes in cluster 9. **d**, Heatmap of genes within clusters 9. Genes and clusters with higher expression in a specific DC type compared to others are shown in red, while genes with lower expression are shown in blue. Data are based on 3 to 5 individual donors per condition. Statistical significance was determined by one-way ANOVA; \**p* < 0.05, \*\**p* < 0.01, \*\*\**p* < 0.001.

**Extended Data Fig. 2:**
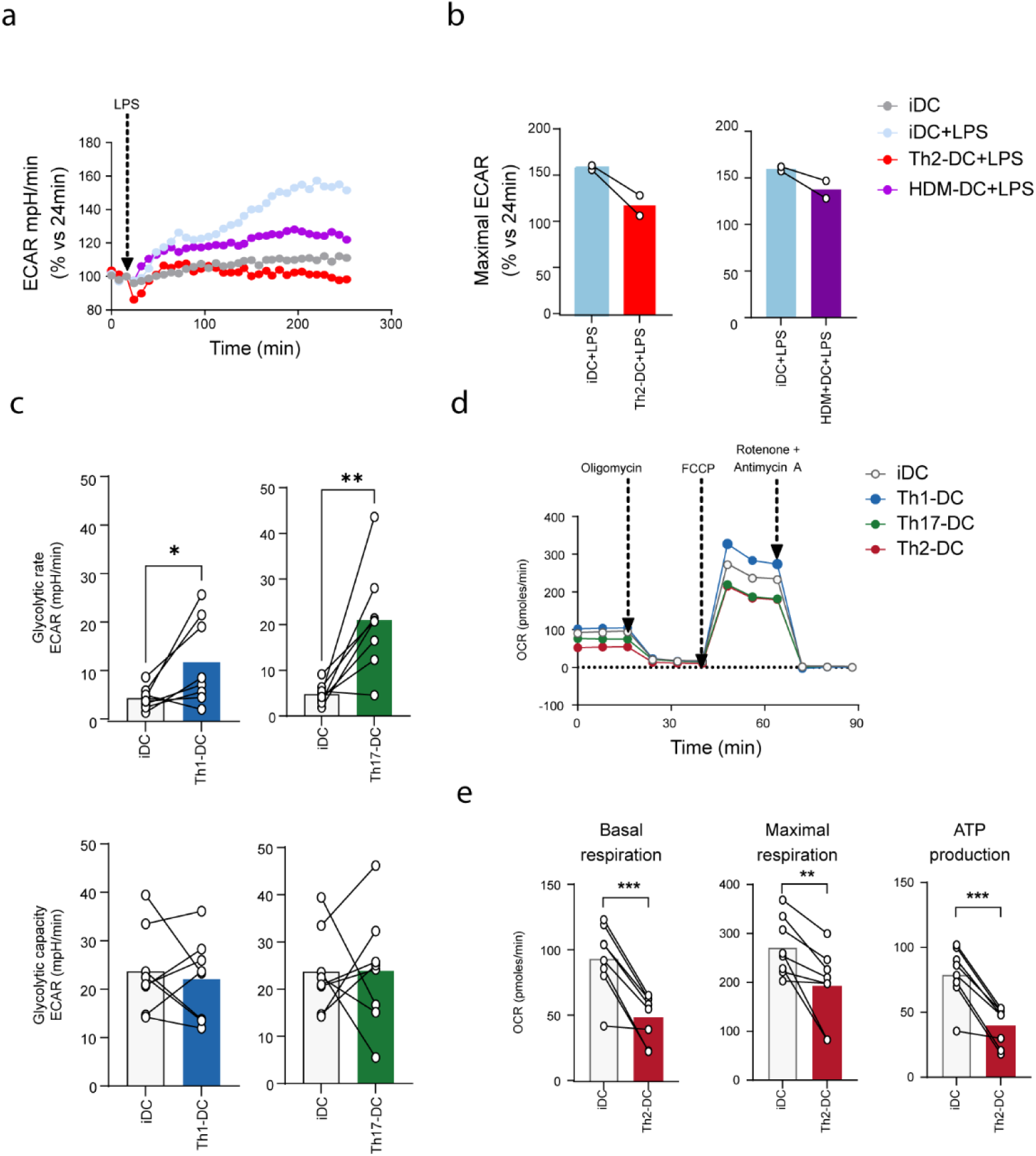
Metabolic characterization of Th1-, Th17- and Th2-priming human moDCs. **a,** Realtime analysis of ECAR following stimulation with LPS in differently pre-conditioned DCs by Seahorse extracellular flux analysis. **b,** Quantification of the data from (**a**). % change in ECAR 180 minutes after LPS injection. **c,** Conditioned DCs were subjected to a glycolytic stress test as described in Figure 3, to determine glycolytic rates and capacity in a Seahorse extracellular flux analyzer. Glycolytic rate is defined as the change in measurement before and after glucose injection. Glycolytic capacity is calculated as the difference between the baseline (pre-glucose injection) and the measurement following oligomycin injection. **d,** Conditioned DCs were subjected to a mitochondrial stress test, in which real-time Oxygen consumption rate (OCR) following oligomycin, FCCP and rotenone/antimycin A were measured by Seahorse extracellular flux analysis. **e,** Basal, maximal and ATP synthesis-coupled respiration were determined in conditioned DCs. Data is shown as mean; each symbol represents a single donor. Statistical significance was determined by paired student’s T test. \**p* < 0.05, \*\**p* < 0.01, \*\*\**p* < 0.001.

**Extended Data Fig. 3:**
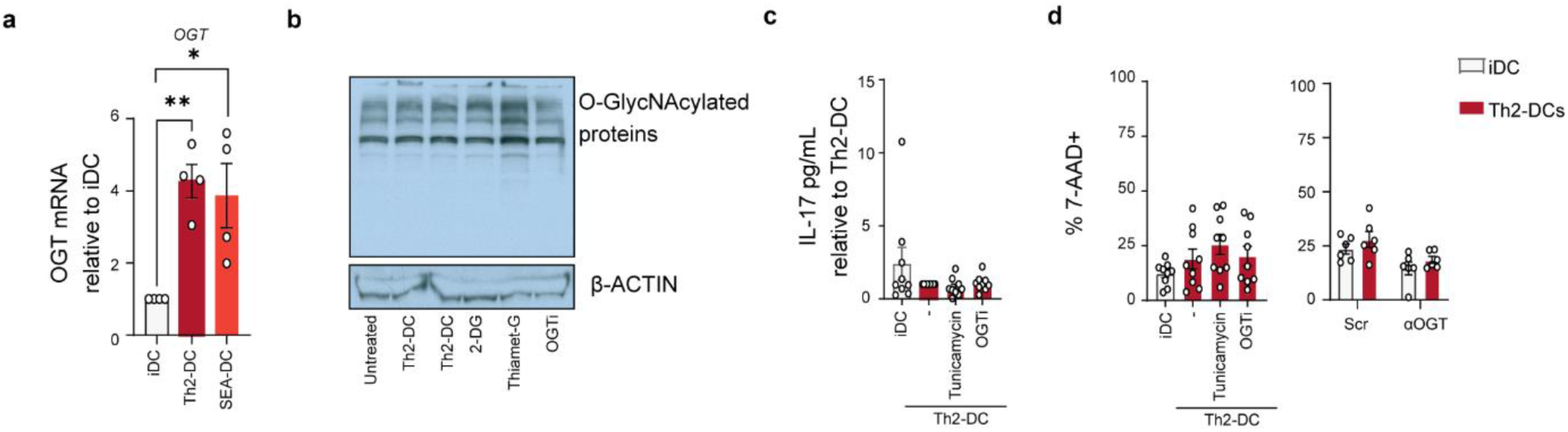
Characterization of O-GlcNAcylation in Th2-priming human moDCs. **a**, mRNA expression of *OGT* in human moDCs stimulated for 24h with either omega-1 (Th2-DC in red) or *Schistosoma mansoni* soluble egg antigens (SEA-DC) as determined by RT qPCR. **b**, Western blot-based analysis of overall protein O-GlcNAcylation on DCs treated with 2-deoxyglucose (2-DG) – an inhibitor of glycolysis, Thiamet G – an inhibitor of O-GlcNAcase and positive control, and ST045849 (OGTi) - an inhibitor of O-GlcNAc transferase. β-Actin was taken along as housekeeping protein. **c**, conditioned-DCs were cocultured with memory Th cells and 5 days after DC-T cell coculture and IL-17 concentrations were determined by ELISA. Relative concentrations are shown. **d**, Frequencies of dead cells in conditioned DCs as determined by 7AAD staining using flow cytometry. Data points represent independent experiments of individual donors. Bars represent mean ± SEM. **a**, Statistical significance was determined using one-way ANOVA with Dunnet post-hoc test. *p < 0.05, **p < 0.01.

**Extended Data Fig. 4:**
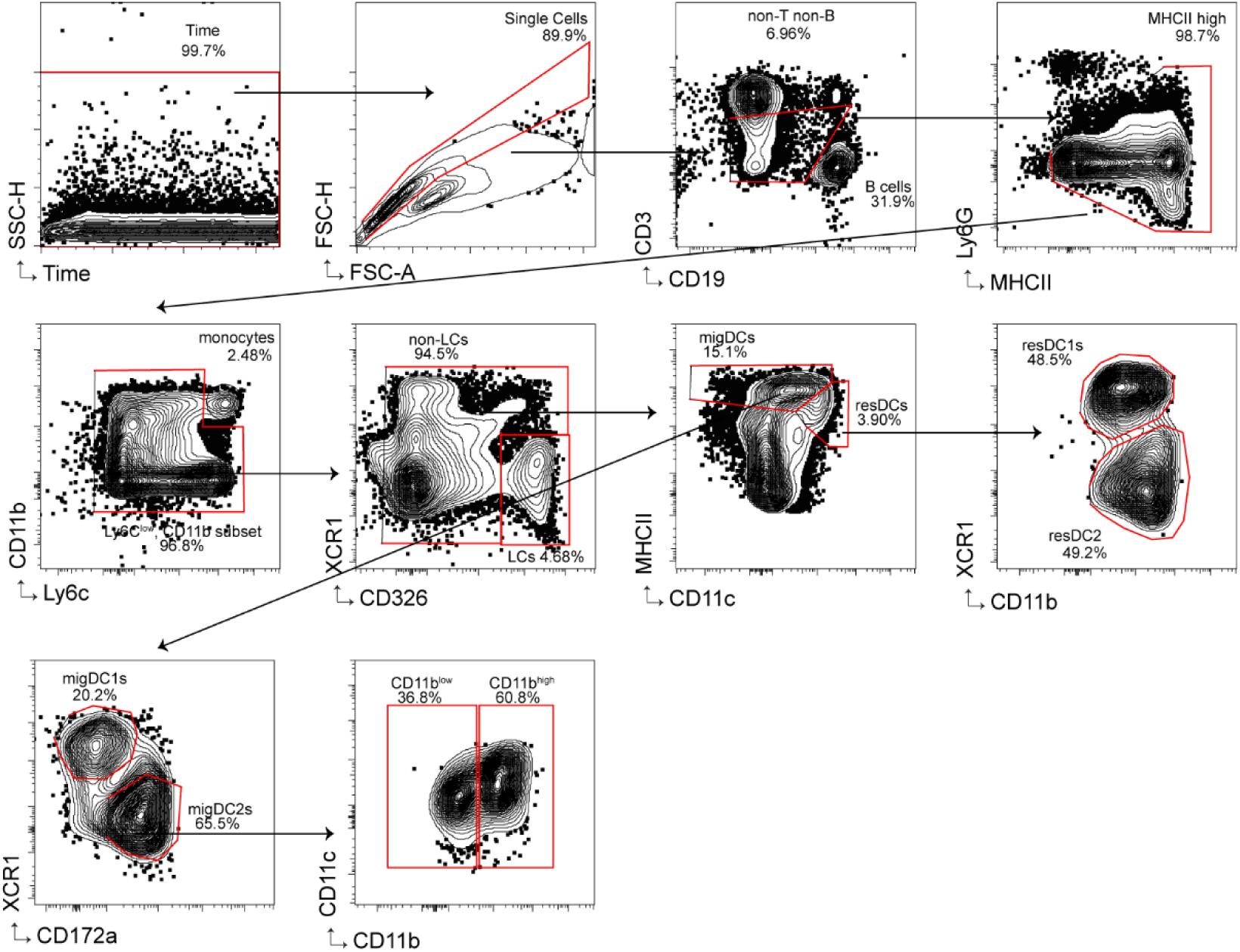
Flow cytometric gating strategy for DC population in skin draining LNs. Representative flow cytometry plots of sequential gating on live CD45⁺ cells, followed by exclusion of lineage-positive cells (CD3, CD19, Ly6G, Ly6C). CD326⁺ Langerhans cells and MHCII^+^CD11c^+^ resident and migratory DCs were defined based on surface marker expression as XCR1⁺ cDC1s, CD172a⁺ cDC2s, and CD11b^low^ double-negative cDC2s (DNDCs).

**Extended Data Fig. 5:**
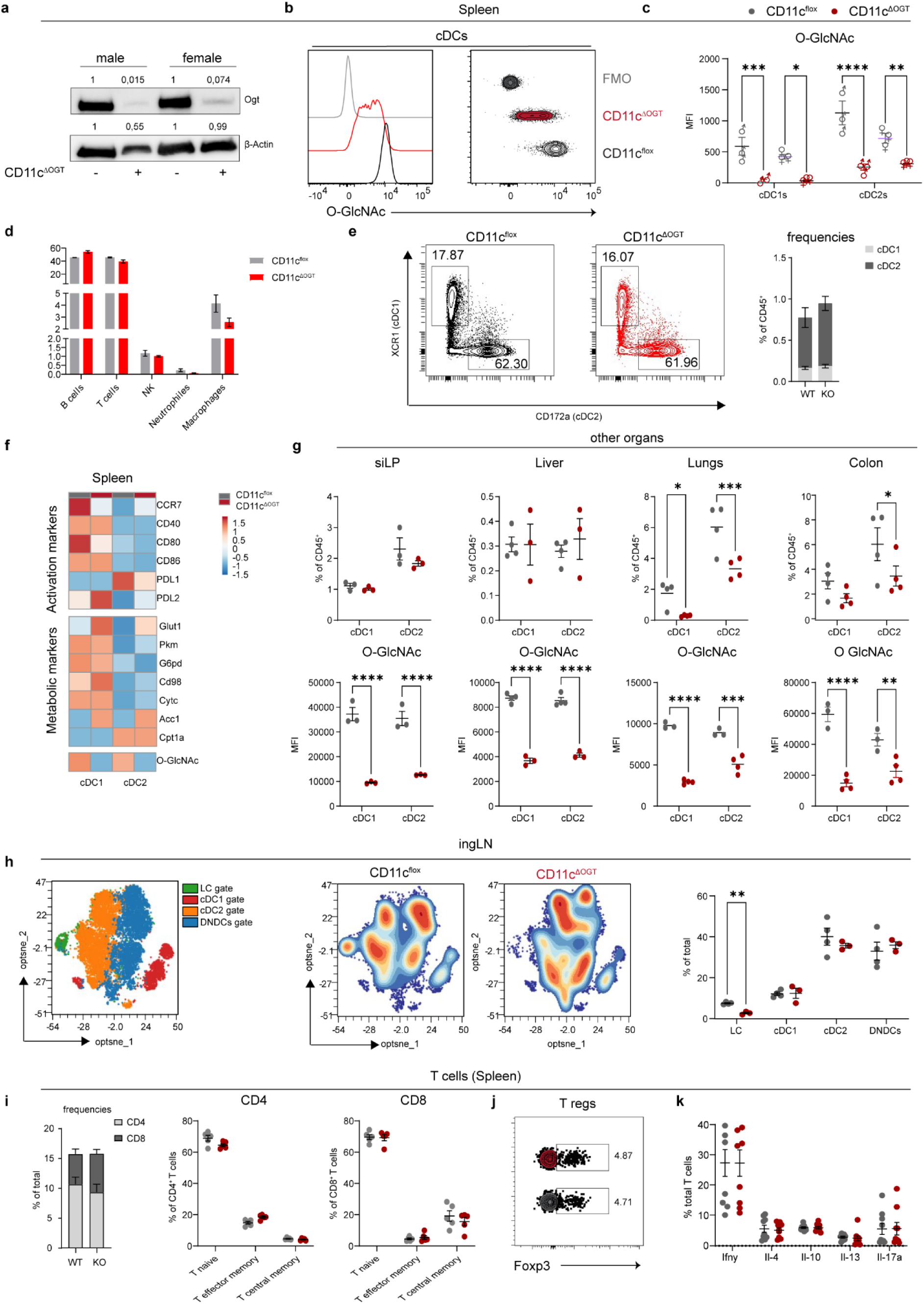
Immunological characterization of naive CD11c^Δ*Ogt*^ mice. **a,** Western blot analysis of Ogt protein expression in FACS sorted CD11c^+^ splenic DCs from CD11c^flox^ and CD11c^Δ*Ogt*^ mice. **b,c,** Flow cytometry based analysis of global levels of protein O-GlcNAcylation in DC subsets isolated from spleens of male and female mice. **b**, representative histograms of O-GlcNAc staining, which is quantified in (**c**). **d**, Frequencies of major immune cell populations and **e**, conventional DCs (cDC1, XCR1⁺; cDC2, CD172a⁺) in spleen. **f**, Heatmap of log2 fold relative differences in expression of activation and metabolic markers in splenic DCs from naïve CD11c^flox^ and CD11c^Δ*Ogt*^ mice. Data are based on average of MFI of indicated markers from 3-4 mice per group. **g,** Frequencies of cDCs and levels of global protein O-GlcNAcylation from the small-intestinal lamina propria (siLP), liver, lung, and colon of naïve CD11c^flox^ and CD11c^Δ*Ogt*^ mice. **h**, Opt-SNE analysis on migratory DCs from skin-draining IngLNs in which (Left) manually gated (Gating in Extended Data Figure 4) populations of Langerhans cells (LC), cDC1, CD11b^+^ cDC2 and (CD24^-^CD11b^-^) DNDCs are indicated or (middle) on which contour plots of naïve CD11c^flox^ and CD11c^Δ*Ogt*^ mice are overlaid. Right: Frequencies of migratory DC subsets. **i**, Frequencies of splenic total, naive, effector and central memory CD4⁺ and CD8⁺ T cells based on CD44 and CD62L expression and of **j**, Foxp3^+^ Tregs in naïve CD11c^flox^ and CD11c^Δ*Ogt*^ mice. **k**, Cytokine production by splenic T cells from naïve CD11c^flox^ and CD11c^Δ*Ogt*^ mice following PMA/ionomycin restimulation. Datapoints represent individual mice. One representative of (**b-e**) 3 or (**f-j**) 2 independent experiments is shown or (**k**) are a pool of two independent experiments. Data are shown as mean ± SEM. Statistical significance was determined by unpaired Student’s t-test and two-way ANOVA with Tukey post-hoc test. \**p* < 0.05, \*\**p* < 0.01, \*\*\**p* < 0.001, \*\*\*\**p* < 0.0001.

**Extended Data Fig. 6:**
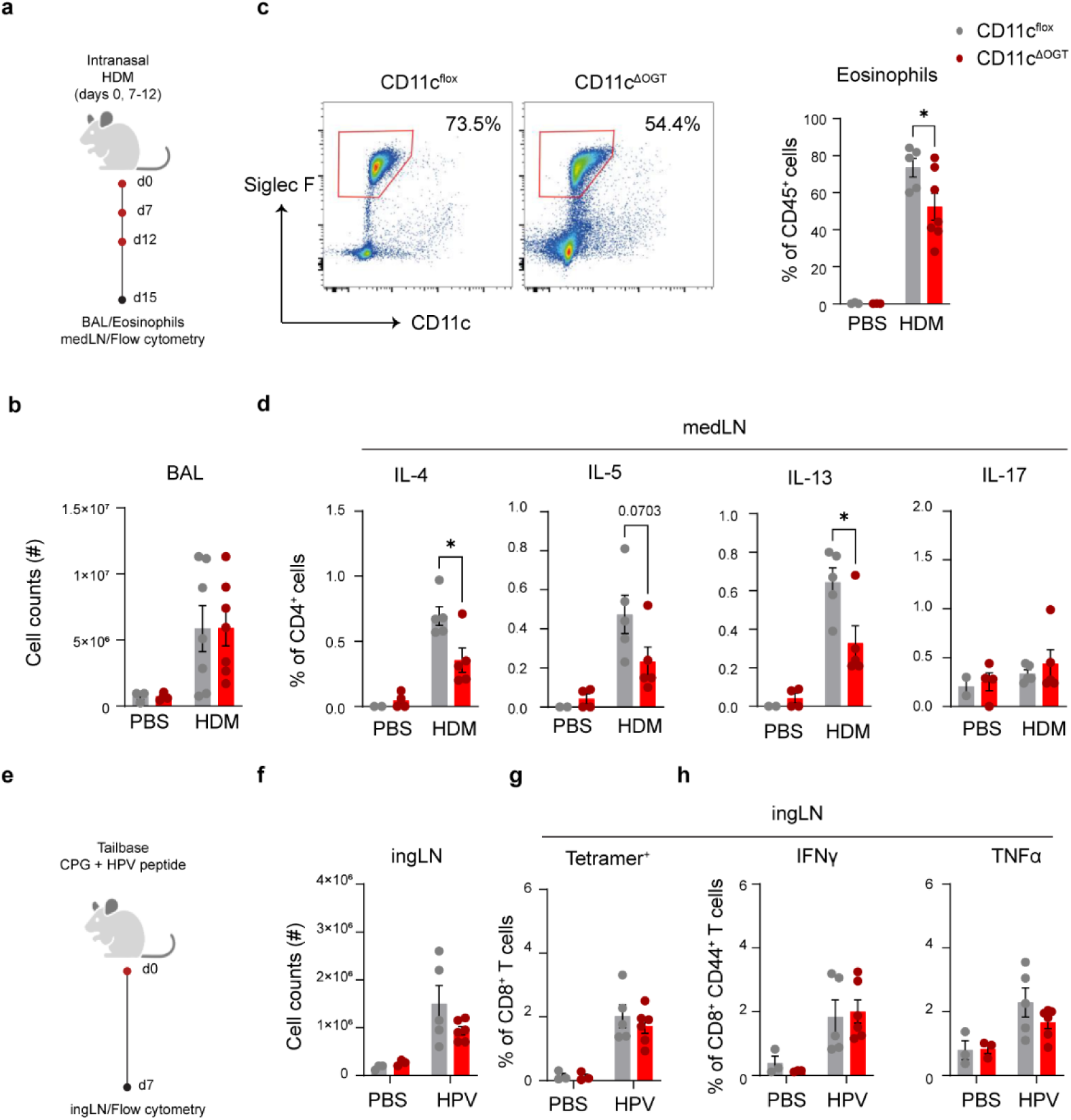
DC O-GlcNAcylation is required for Th2 priming in house-dust-mite (HDM)–induced asthma but not for CD8^+^ T cell priming *in vivo*. **a,** Experimental model of intranasal HDM administration with sensitization on day 0 and allergen challenge days on 7–12. Bronchoalveolar lavage (BAL) fluid and mediastinal lymph nodes (medLN) were collected on day 15. **b,** Number of cells in BAL. **c**, Flow-cytometric quantification of eosinophils in BAL**. d,** Cytokine expression in CD4⁺ T cells following 6h PMA/ionomycin restimulation of medLN cells *ex vivo*. **e**, Experimental model of HPV peptide plus CpG immunization at the tail base. Inguinal lymph nodes (ingLN) were harvested 7 days post-immunization. **f**, IngLN Cellularity. **g,** E7-Tetramer staining and **h,** IFN-γ and TNF expression in CD8⁺ T cells after 6-h PMA/ionomycin restimulation of IngLN cells *ex vivo*. Datapoints represent individual mice from one experiment (**a-d**) or one representative experiment out of two (**e-g**). Data are shown as mean ± SEM. Statistical significance was determined by two-way ANOVA with Tukey post-hoc test. \**p* < 0.05.

**Extended Data Fig. 7:**
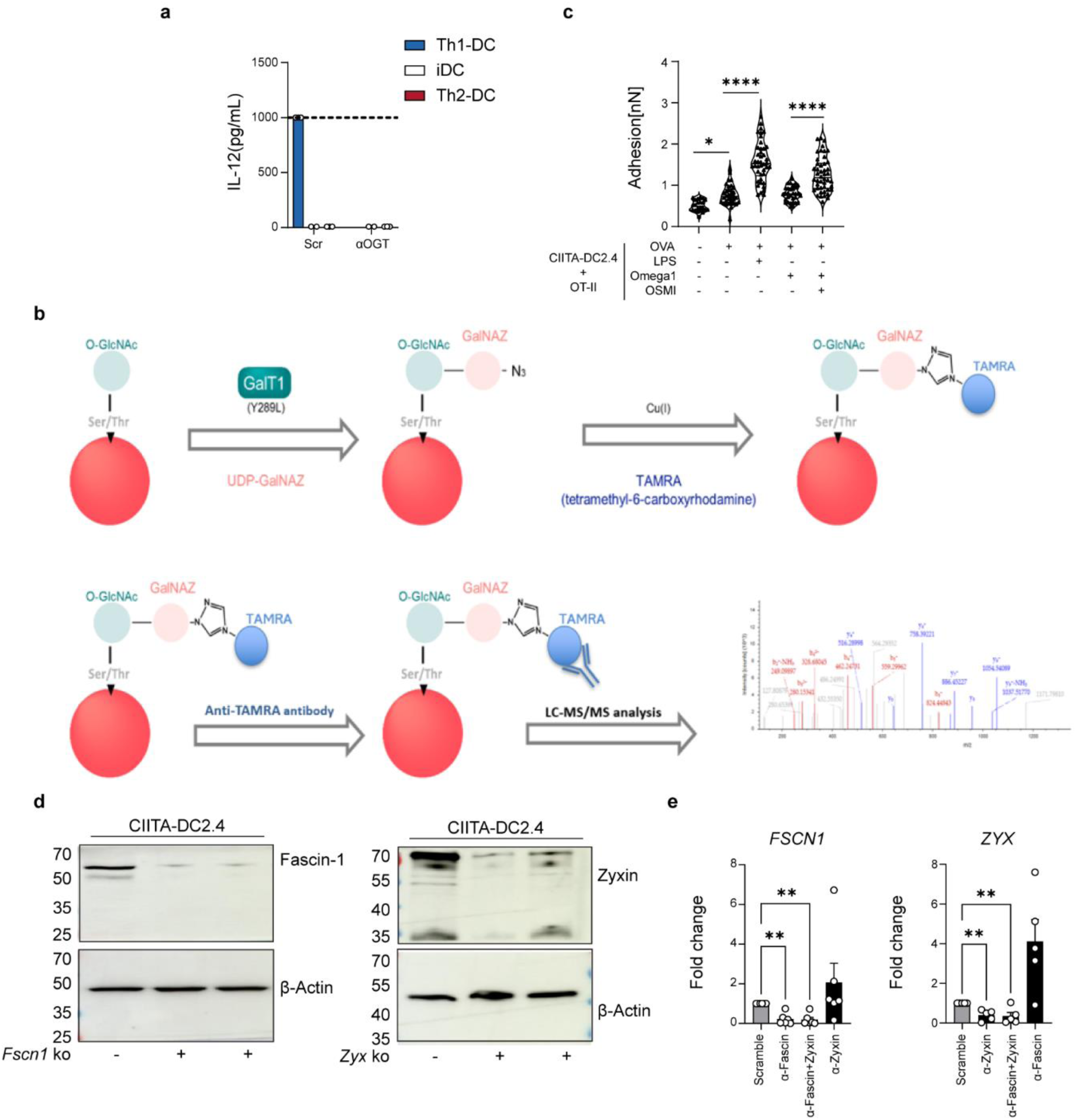
Mapping O-GlcNAcylation targets and targeting of Zyxin and Fascin. **a**, Differentiating human moDCs were transduced with siRNA against O-GlcNAc transferase (*OGT*) on day 4. Scrambled RNA was used as a control. At day 6, DCs were stimulated with either LPS+Poly(I:C) (Th1-DC in light blue), omega-1 (Th2-DC in red) for 24 hours or untreated (iDC in grey). Supernatants were then collected for determination of IL-12p70 secretion. **b**, Proteomic determination of global O-GlcNAcylome. O-GlcNAcylated proteins in stimulated DCs were first enzymatically tagged with a GalNAZ residue, followed by click-chemistry based tagging with a fluorescent TAMRA label. Anti-TAMRA-immunoprecipitated proteins were identified via LC-MS/MS. **c**, DC2.4 cells were stimulated with LPS or omega1 with or without pre-incubation with OGT inhibitor OSMI prior to stimulation. The adhesion properties between OTII cells and different treated DC2.4 cells have been quantified. **d**, Western blot for Fascin-1 and Zyxin on DC2.4 cells in which Fascin-1 and Zyxin were knocked out by CRISPR/Cas9. **e**, mRNA expression of *FSCN1* and *ZYX* was determined by RT qPCR after siRNA-mediated silencing in human moDCs. Data are shown as mean ± SEM. Statistical significance was determined by one-way ANOVA with Tukey post-hoc test. *p < 0.05, \*\**p* < 0.01, \*\*\*\**p* < 0.0001.

